# A unifying structural and functional model of the coronavirus replication organelle: tracking down RNA synthesis

**DOI:** 10.1101/2020.03.24.005298

**Authors:** Eric J. Snijder, Ronald W.A.L. Limpens, Adriaan H. de Wilde, Anja W. M. de Jong, Jessika C. Zevenhoven-Dobbe, Helena J. Maier, Frank F.G.A. Faas, Abraham J. Koster, Montserrat Bárcena

## Abstract

Zoonotic coronavirus (CoV) infections, like those responsible for the current SARS-CoV-2 epidemic, cause grave international public health concern. In infected cells, the CoV RNA-synthesizing machinery associates with modified endoplasmic reticulum membranes that are transformed into the viral replication organelle (RO). While double-membrane vesicles (DMVs) appear to be a *pan*-coronavirus RO element, studies to date describe an assortment of additional coronavirus-induced membrane structures. Despite much speculation, it remains unclear which RO element(s) accommodate viral RNA synthesis. Here we provide detailed 2D and 3D analyses of CoV ROs and show that diverse CoVs essentially induce the same membrane modifications, including the small open double-membrane spherules (DMSs) previously thought to be restricted to gamma- and delta-CoV infections and proposed as sites of replication. Metabolic labelling of newly-synthesized viral RNA followed by quantitative EM autoradiography revealed abundant viral RNA synthesis associated with DMVs in cells infected with the beta-CoVs MERS-CoV and SARS-CoV, and the gamma-CoV infectious bronchitis virus. RNA synthesis could not be linked to DMSs or any other cellular or virus-induced structure. Our results provide a unifying model of the CoV RO and clearly establish DMVs as the central hub for viral RNA synthesis and a potential drug target in coronavirus infection.

## Introduction

The RNA synthesis of all positive-stranded RNA (+RNA) viruses of eukaryotes occurs in the cytoplasm of the host cell, in conjunction with modified endomembranes that are often referred to as viral replication organelles (ROs) [1–3]. ROs are generally believed to provide tailored platforms that facilitate viral replication by concentrating relevant factors and spatially organizing distinct steps in the viral cycle. Additionally, ROs may contribute to the evasion of cellular innate immune defences that detect viral RNA (vRNA) [4].

Two main RO prototypes have been discriminated: small spherular invaginations and large(r) vesiculotubular clusters consisting of single- and/or double-membrane structures, to which viral replicative proteins and specific host factors can be recruited. The formation of invaginations can occur at the membrane of various organelles, including endoplasmic reticulum (ER), endolysosomes, and mitochondria [5–9]. The lumen of the resulting micro-compartment is connected with the cytosol by a ‘neck-like’ channel that can mediate transport of metabolites and export of newly made positive-sense vRNAs to the cytosol for translation and packaging. In general, the morphological and functional characterization of ROs of the second, vesiculotubular type is lagging behind. Such structures, which always include double-membrane vesicles (DMVs), commonly derive from membranes of the secretory pathway and have been found in cells infected with e.g. picornaviruses [10, 11], noroviruses [12], hepatitis C virus (HCV) [13], and different nidoviruses, including the arterivirus and coronavirus (CoV) families [14–18].

The first electron tomography analysis of a coronavirus-induced RO, that of the severe acute respiratory syndrome-CoV [15] (SARS-CoV, genus *Betacoronavirus*), raised a variety of functional considerations. Intriguingly, while double-stranded (ds) RNA, a presumed intermediate and marker for vRNA synthesis [19], was found inside the virus-induced DMVs, these lacked visible connections to the cytosol [15]. Viral RNA synthesis inside fully-closed DMVs would pose the conundrum of how metabolites and newly made genomic and subgenomic mRNAs could be transported across the double-lipid bilayer. Nevertheless, experimental evidence for the location of vRNA synthesis within the RO was lacking. Importantly, dsRNA is not a *bona fide* marker for vRNA synthesis as it may no longer be associated with the active enzymatic replication complexes in which most of the 16 viral non-structural proteins (nsps) come together. Thus, the possibility of vRNA synthesis taking place in alternative locations, like the convoluted membranes (CM) that are also prominent elements of the beta-CoV RO [15, 20, 21], was entirely possible and started to attract attention. Notably, DMVs can be also formed in the absence of vRNA synthesis by expression of key transmembrane nsps [22–26]. Moreover, several studies suggested a lack of direct correlation between the number of DMVs and the level of CoV replication in the infected cell [27, 28].

The interpretation of the CoV RO structure and function was further compounded by the discovery of different RO elements, never reported in beta-CoV infections, that set apart other distantly-related CoVs genera. In particular, zippered ER (instead of CM) and double-membrane spherules (DMSs) were detected for the avian gamma-CoV infectious bronchitis virus (IBV) [29] and, recently, for the porcine deltacoronavirus [30]. The size and topology of these DMSs, invaginations in the zippered ER, were remarkably similar to those of the spherular invaginations induced by other +RNA viruses and, consequently, DMSs were suggested to be sites of vRNA synthesis [29, 31].

In this work, we provide an in-depth analysis of the structure and function of the RO induced by members of different CoV genera, with a special focus on beta-CoVs like SARS-CoV and MERS-CoV (Middle East respiratory syndrome-CoV) [32, 33]. While the SARS-CoV outbreak was contained in 2003, MERS-CoV continues to pose a serious zoonotic threat to human health since 2012 (http://www.who.int/emergencies/mers-cov/en/). Very recently, a previously unknown beta-CoV (2019-nCoV, recently renamed into SARS-CoV-2) has emerged in China [34] and is causing major public health concerns worldwide. The 2019-nCoV genome sequence is 79.5% identical to that of SARS-CoV [35], yielding 86% overall nsp sequence identity and suggesting strong functional similarities in the replication of both viruses.

Our observations on the 3D morphology of the MERS-CoV RO were compared with data from cells infected with other alpha-, beta-, and gamma-CoVs. This comparative analysis made it clear that all these CoV induce essentially the same membrane structures, including DMSs. Metabolic labelling of newly-synthesized vRNA was used to determine the site(s) of vRNA synthesis within the CoV RO. To this end, we used a radiolabeled nucleoside ([^3^H]uridine) and applied the classic and highly-sensitive technique of EM autoradiography [36, 37] in combination with advanced quantitative analysis tools. This approach revealed that DMVs are the primary site of CoV RNA synthesis, with neither DMSs nor CM or zippered ER being labelled to a significant extent. Our study provides a comprehensive and unifying model of the CoV RO structure. It also returns DMVs to centre stage as the hub of CoV RNA synthesis and a potential antiviral drug target.

## Results

### MERS-CoV induces a membrane network of modified membranes that contains double-membrane spherules (DMSs)

We first set out to analyse the ultrastructure of MERS-CoV-infected Huh7 cells under sample preparation conditions favourable for autoradiography (see Materials and Methods) (Fig 1, S1 Video). Strikingly, in addition to the DMVs and CM that are well established hallmarks of beta-CoV infections, the presence of small spherules, occasionally in large numbers, was readily apparent (Fig 1A and 1B). These spherules were notably similar to the DMSs previously described for the gamma-CoV IBV [29]. Their remarkably regular size of ~80 nm (average diameter 79.8 ± 2.5 nm, n=58), a delimiting double membrane and their electron-dense content, made these spherules clearly distinct from other structures, including progeny virions, which had comparable diameter (Fig 1C and 1D). The double-membrane spherules (DMSs) generated during IBV infection were previously described as invaginations of the zippered ER that remain open to the cytosol [29]. In MERS-CoV-infected cells, the DMSs were connected to the CM from which they seemed to derive (Fig 1E). Clear openings to the cytosol could not be detected for the large majority (~80%, n=54) of the fully reconstructed DMSs, which suggests that the original invagination may eventually transform into a sealed compartment. This type of apparently closed DMSs were also present, though in a lower proportion (~50%, n=39), in IBV-infected cell samples processed in an identical manner (S1 Fig).

**Fig 1.**
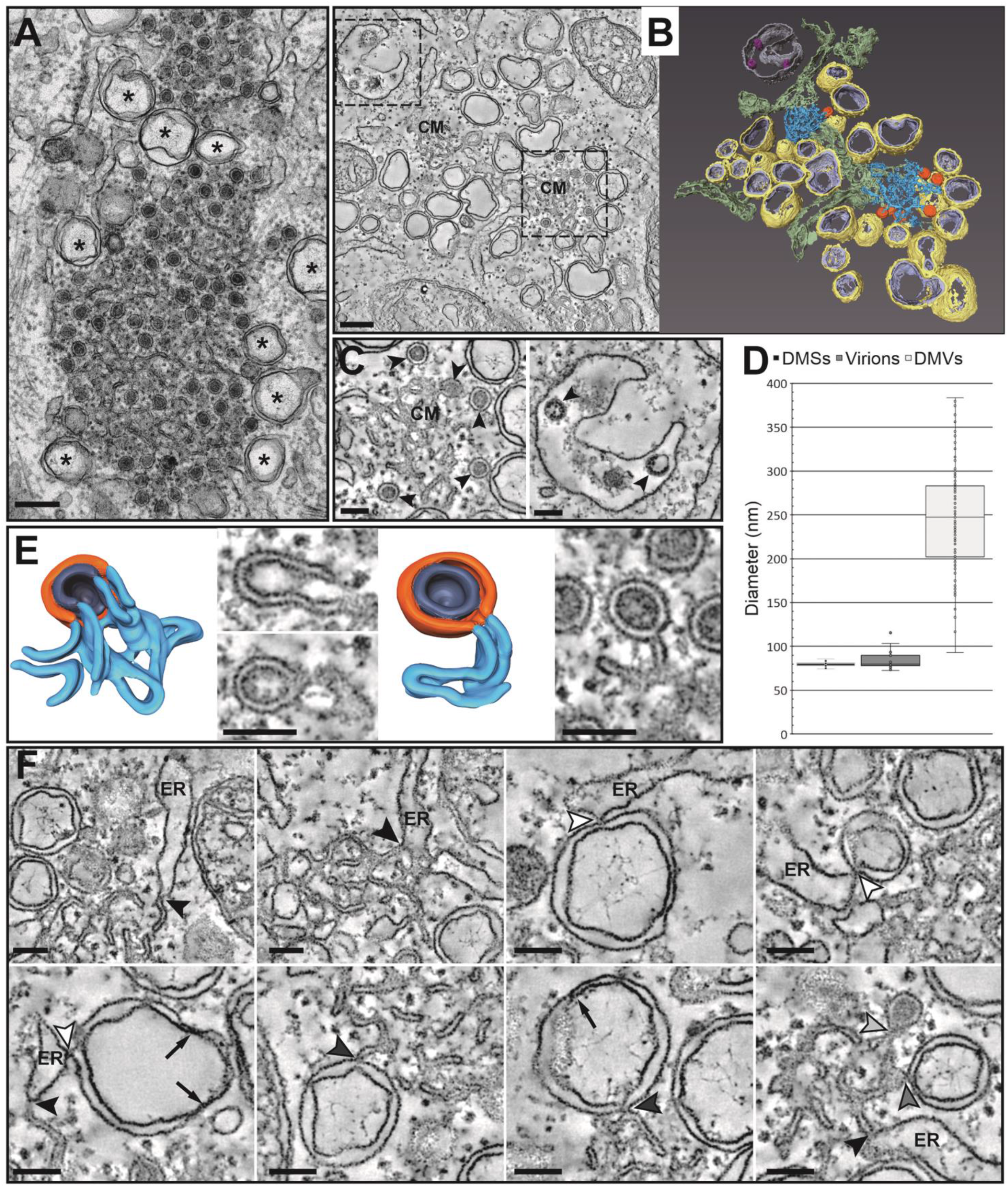
Membrane structures induced by MERS-CoV infection. Electron microscopy analysis of Huh7 cells infected with MERS-CoV (MOI 5, 12 hpi). (A) Electron micrograph of an area with abundant double-membrane spherules (DMSs). DMVs (asterisks) are interspersed and surrounding the DMS cluster. (B) Slice through a tomogram (left) and corresponding surface-rendered model (right) of a representative area containing the different types of MERS-CoV-induced membrane modifications: DMSs (orange), convoluted membranes (CM, blue) and DMVs (yellow and lilac, outer and inner membranes). The model also highlights ER membranes (green) and a vesicle (silver) containing new virions (pink). (See also S1 Video). (C) Comparison of DMSs and virions (arrowheads in left and right panels, respectively) in enlarged views of tomographic slices from the regions boxed in (B). The DMSs are similar in size but distinct in appearance from newly-formed MERS-CoV particles. (D) Whisker plots of the size distribution of DMSs (n=58), virions (n=28) and DMVs (n=109), as measured from the tomograms. DMSs and virions have a comparable size (median diameter, 80 nm), while the median diameter of the DMVs is 247 nm. (E) Models and tomographic slices through an open (left) and closed (right) DMS. Both types of DMSs are connected with the CM. In open DMSs, both the inner and outer membranes (dark blue and orange, respectively) are continuous with CM. Two ~8-nm apart slices through the reconstruction are shown. For closed DMSs only the outer membrane is connected to CM, while the inner membrane seems to define a closed compartment. (F) Gallery of tomographic slices highlighting membrane connections between different elements of the MERS-CoV RO, and of these with the ER. These include CM-ER (black arrowheads), DMV-ER (white arrowheads), CM-DMV (dark grey arrowheads), and CM-DMS (light grey arrowhead) connections. Scale bars, 250 nm (A, B), and 100 nm (C -F).

Our data suggests a functional analogy between zippered ER and CM, as both structures appear to provide membranes for DMS formation. In fact, both bore a striking resemblance under the same sample preparation conditions: MERS-CoV-induced CM noticeably consisted of zippered smooth membranes that were connected to the rough ER and branched and curved in intricate arrangements. Although unbranched zippered ER was most common in IBV-infected cells, as described [29], we also detected zippered ER morphologically closer to CM (S1B Fig). These observations argue for zippered ER and CM representing alternative configurations of essentially the same virus-induced RO element.

The 3D architecture of MERS-CoV-induced RO aligned with previous observations for other CoV [15, 29]. No clear openings connecting the interior of the DMVs and the cytosol could be detected. All three types of MERS-CoV-induced membrane modifications appeared to be interconnected, either directly or indirectly through the ER. While DMSs were connected to CM, and CM to ER, ER membranes were often continuous with DMVs (Fig 1F, arrowheads). Therefore, like other CoVs, MERS-CoV infection appears to induce a network of largely interconnected modified ER membranes that, as a whole, can be considered the CoV RO.

### Diverse coronaviruses across different genera induce the same RO elements

Intriguingly, DMSs had never been reported for beta-CoV infections in previous characterizations, including ours, that used different sample preparation conditions and/or different cell lines. This prompted us to revisit those samples for a closer examination. A targeted search for DMSs in MERS-CoV-infected Vero cells [21] and SARS-CoV-infected Vero E6 cells [15] readily revealed similar DMSs, embedded and somewhat concealed in CM with a denser and more tangled appearance that can be attributed to sample preparation differences (S2 Fig, compare to Fig 1 and Fig 2A). To further explore whether these observations could be extended to other coronaviruses, we analysed a third beta-CoV (murine hepatitis virus, MHV) as well as a member of the genus *Alphacoronavirus* (human coronavirus 229E, HCoV-229E). Both in MHV-infected 17Cl1 cells and in HCoV-229E-infected Huh7 cells, virus-induced DMSs could be detected, with their characteristic size, appearance, and spatial association with CM (Fig 2B and 2C). These results demonstrate that virus-induced DMSs are not exclusive to some CoV genera or

**Fig 2.**
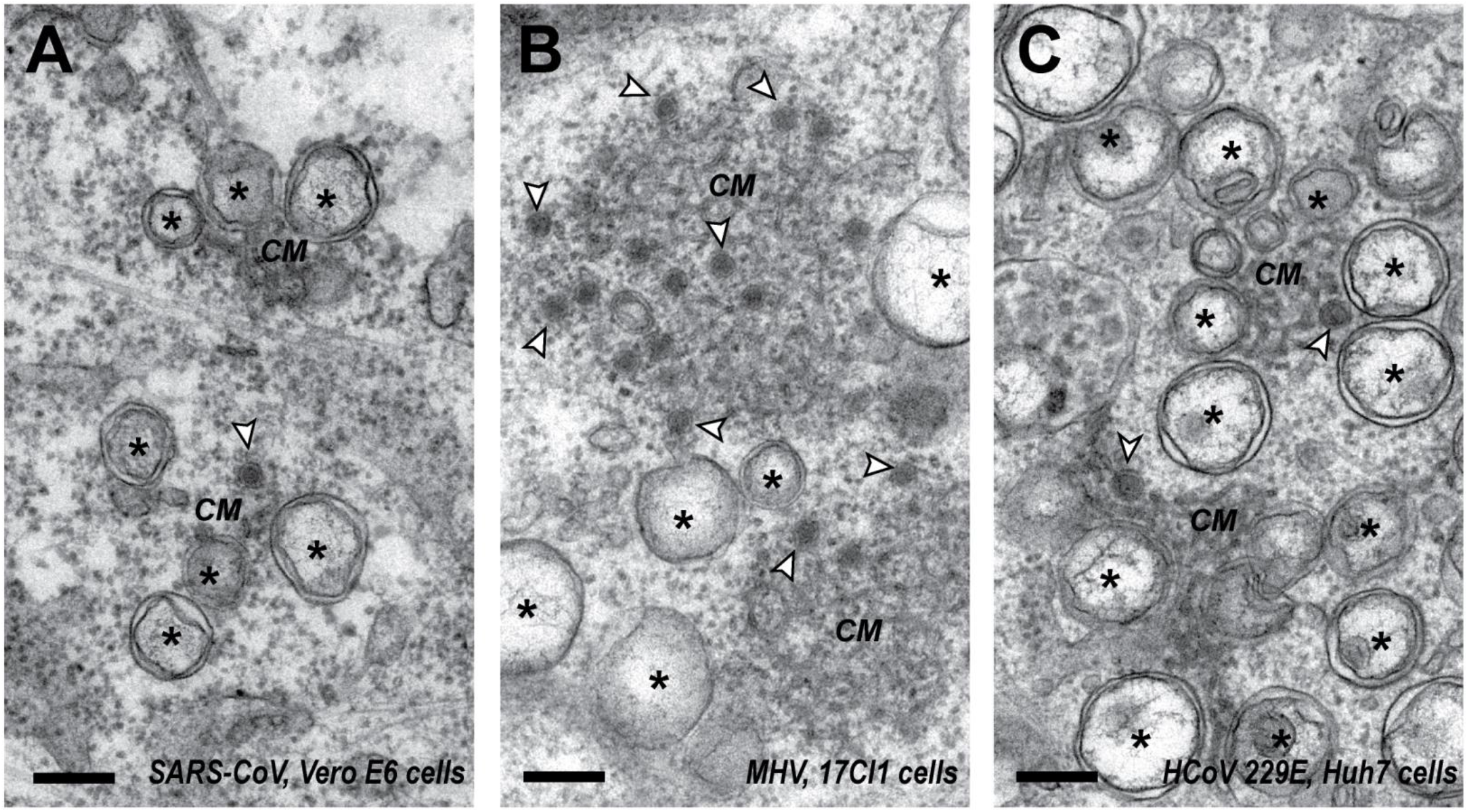
DMSs are induced by diverse beta- and alpha-CoVs. 2D-EM images from 100-nm thick sections of different mammal cells infected with (A) SARS-CoV (MOI 10, 9 hpi), (B) MHV (MOI 10, 8 hpi) and (C) HCoV-229E (MOI 5, 24 hpi). Both beta-CoVs (A,B) and the alpha-CoV (C) induce membrane modifications that include not only DMVs (asterisks) and CM, but also DMSs (white arrowheads). Scale bars, 250 nm.

### Coronavirus RNA synthesis is confined to RO regions

To investigate the subcellular localization of vRNA synthesis in cells infected with different CoVs, we metabolically labelled newly-synthesized vRNA by arresting cellular transcription with actinomycin D and using a radiolabelled nucleoside precursor ([5-^3^H]uridine) for subsequent detection by EM autoradiography [36, 37] (see S1 Appendix). A key advantage of this approach over the use of modified precursors (e.g. Br-uridine), is that detection of the label does not rely on immunolabelling. This makes autoradiography compatible with high-contrast EM sample preparation protocols that provide excellent morphology at the price of epitope integrity. Moreover, as the signal derives from radioactive disintegrations, EM autoradiography is a very sensitive technique, which in principle should allow for short labelling pulses, essential to minimize the chance of migration of labelled vRNA products from their site of synthesis. Nevertheless, the pulse should be long enough to enable the internalization of the tritiated uridine, its conversion into ^3^H-UTP in the cell, and its incorporation into vRNA. In order to explore the practical limits of the approach, we tested different labelling pulses in Vero E6 cells infected with SARS-CoV, measuring the amount of radioactive label incorporated into RNA (Fig 3A). While minimal labelling occurred within the first 10 minutes, a sharp increase in signal was observed between 10 and 20 min after label administration. Consequently, only samples labelled for 20 min or longer were prepared and processed for EM autoradiography analysis.

**Fig. 3.**
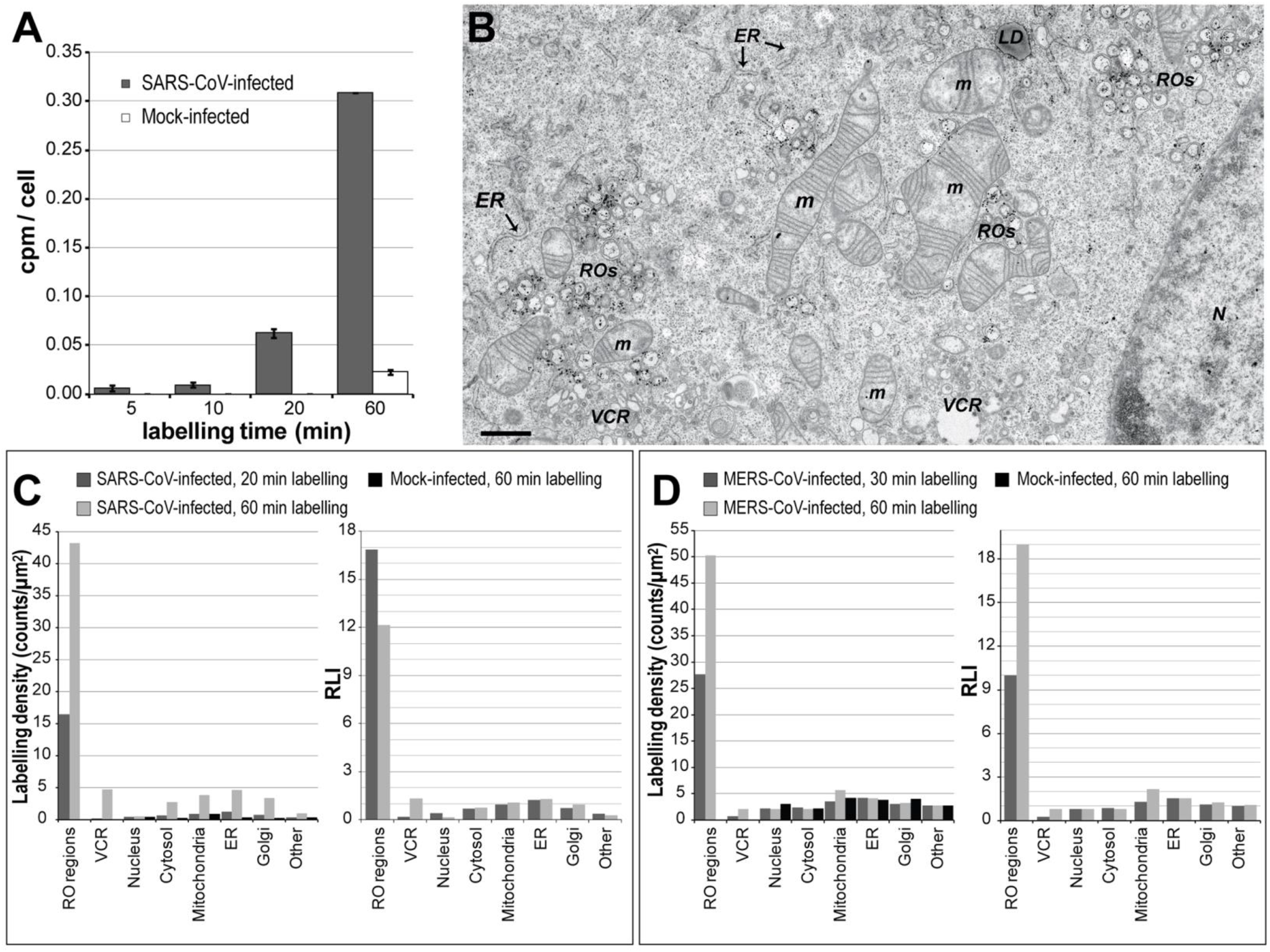
CoV RNA synthesis is confined to RO regions. Newly-synthesized vRNA was metabolically labelled by providing tritiated uridine to CoV-infected cells pre-treated with actinomycin D to limit host transcription. (A) Analysis of the amount of radioactive label incorporated into RNA as a function of the labelling time in SARS-CoV-infected Vero E6 cells (MOI 10), as measured by scintillation counting on the RNA isolated from the cells. The label was provided simultaneously to all the samples at 6 hpi. (B-D) EM detection by autoradiography. (B) Overview of a SARS-CoV-infected Vero E6 cell (MOI 10) were labelled for 20 min just before fixation at 7 hpi, and then processed for EM autoradiography. Autoradiography grains accumulate in the RO regions. N, nucleus; m, mitochondria; VCR, virion-containing regions; LD, lipid droplets. Scale bar, 1 μm. (C, D) Quantification of the autoradiography signal per subcellular structure (see also S1 Table). Labelling densities and relative labelling indexes (RLI) in different subcellular regions of (C) Vero E6 cells infected with SARS-CoV (MOI 10) or (D) Huh7 cells infected with MERS-CoV (MOI 5). Radioactively-labelled uridine was provided for the indicated periods of time immediately before fixation at 7 hpi and 12 hpi, respectively. Control mock-infected cells are excluded from the RLI plots, as RLI comparisons between conditions require the same number of classes (subcellular regions) and these cells lack ROs and virions.

Abundant autoradiography signal was detected by EM in SARS-CoV-infected cells pulse-labelled for 20 min (Fig 3B), accumulating in the regions that contained virus-induced membrane modifications, which aligned with the idea that these structures are the primary platforms for vRNA synthesis. This largely accepted notion, however, does not formally exclude the possibility vRNA synthesis could also be associated with other cellular membranes, albeit to a lower extent. Such an association with morphologically intact membranes could be important, for example, in the first stages of infection, when the levels of the viral membrane-remodelling proteins are still low.

Importantly, establishing the association of autoradiography signal with specific subcellular structures requires a detailed quantitative analysis, as the autoradiography signal can spread up to a few hundred nanometers from the original radioactive source (see S1 Appendix). This type of analysis was used to compare CoV- and mock-infected cells labelled for different periods of time. To this end, we analysed the autoradiography signal present in hundreds of regions that were randomly picked from large EM mosaic images [38] and calculated labelling densities and relative labelling indexes (RLI) per compartment [39] (see Materials and Methods, S1 Table). The results for SARS-CoV did not show association of vRNA synthesis with any subcellular structure other than ROs (Fig 3C). In infected cells labelled for a short period of time (20 min), the RO labelling densities were one order of magnitude higher than for any other subcellular structure. Even though the dispersion of signal around the radioactive source would inevitably cause ‘signal leakage’ from these active ROs to neighbouring organelles like the ER, none of those alternative locations showed an RLI significantly higher than 1 (RLI≤1 indicates unspecific labelling), and their labelling densities were comparable to those in the control mock-infected cells. An increase in labelling densities in some subcellular regions was observed in infected cells when the labelling time was extended to 60 min. This could be explained by signal leakage in combination with the migration of vRNA from its site of synthesis, possibly towards the site of virus assembly on membranes of the ER-Golgi intermediate compartment (ERGIC) [40–42]. Indeed, virion-containing regions showed the sharpest increase in labelling density when extending the labelling time. Similar results followed from the analysis of MERS-CoV-infected Huh7 cells (Fig 3D), although no clear signs of RNA migration were observed in this case. Taken together, these results suggest that CoV RNA synthesis is restricted to the RO-containing regions of the infected cell.

### DMVs are the primary site of coronaviral RNA synthesis

Our next goal was to determine which elements of the coronaviral RO (DMVs, CM/zippered ER and/or DMSs) are directly involved in vRNA synthesis. A first answer to this question became readily apparent as regions containing DMVs, but not either of the other RO structural elements, were densely labelled in cells infected with MERS-CoV, SARS-CoV or IBV (Fig 4A and 4B, Fig 3B, and S3 Fig, respectively). Interestingly, not all the DMV clusters in a sample, and sometimes even within a cell, appeared equally densely labelled, which suggests that the levels of vRNA synthesis in DMVs are variable and could change in time. The active role of DMVs in vRNA synthesis in MERS-CoV-infected cells was further corroborated by a detailed analysis of the distribution of autoradiography signal around isolated DMVs (Fig 4C). In case of a random distribution (i.e. if the DMVs were not a true radioactive source), the number of autoradiography grains around the DMVs would simply increase with the distance, as the perimeter of the screened area also increases and with it the chances of detecting background signal. The signal around DMVs was clearly not random, showing maximum levels in the proximity of these structures (Fig 4C). Moreover, the distribution normalized by the distance made apparent a maximum around the average radius of the DMVs analysed (133 ± 28 nm, n=36) (Fig 4E), aligning with the idea that vRNA synthesis takes place in membrane-bound enzymatic complexes.

**Fig 4.**
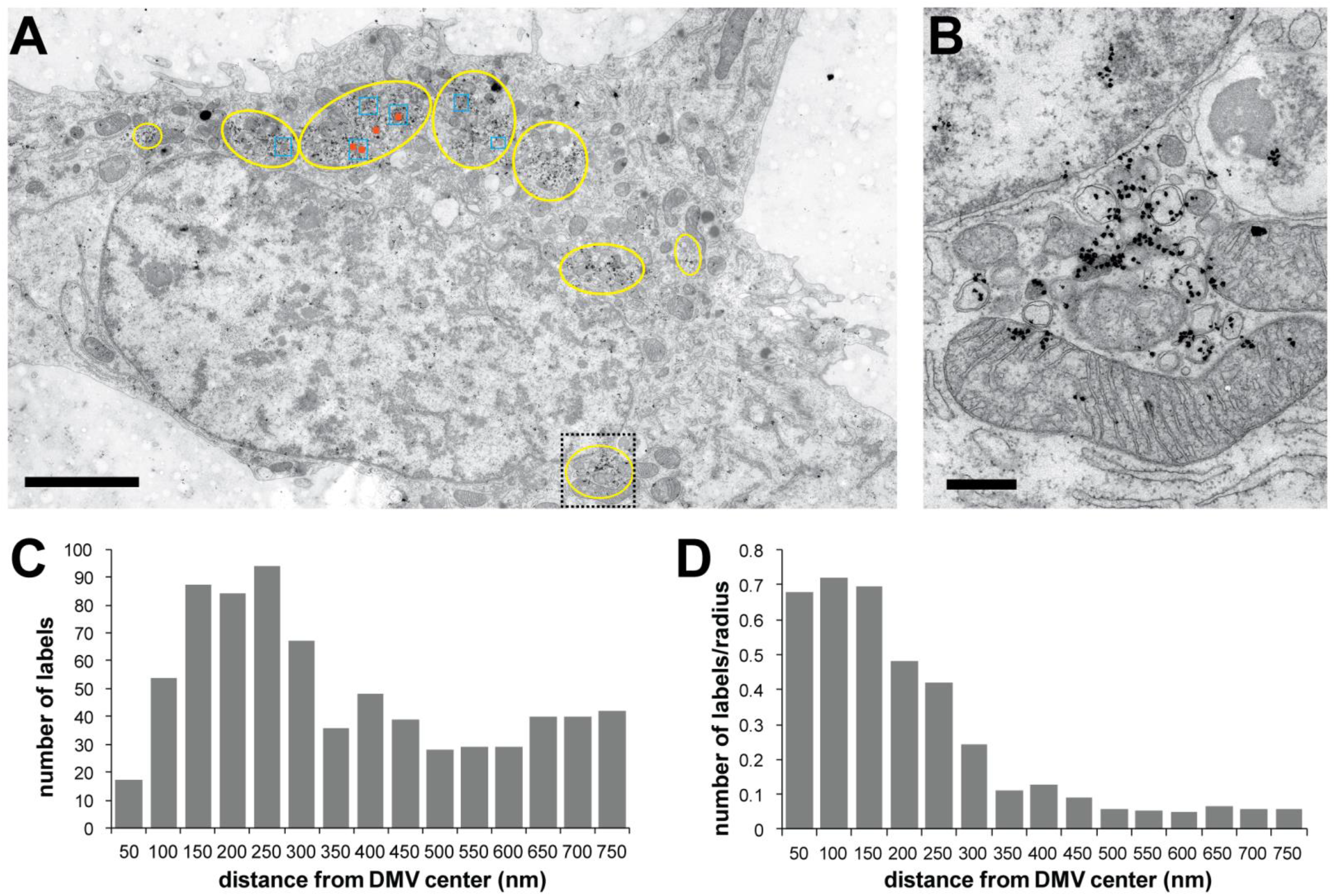
DMVs are sites of vRNA synthesis. Analysis of the association of autoradiography signal with DMVs in MERS-CoV-infected Huh7 cells (MOI 5). The cells were pre-treated with actinomycin D at 10 hpi, and labelled with tritiated uridine for 30 min immediately before fixation (12 hpi). (A) Overview of an infected cell in which regions with different virus-induced modifications are annotated in yellow (DMVs), blue (CM) and orange (DMSs). Several densely-labelled regions containing DMVs, but not the other virus-induced structures are apparent. A close-up of one of these regions (boxed area) is shown in (B). (C, D) Distribution of the autoradiography signal around DMVs (n_DMVs_ = 36, see Material and Methods for selection criteria and details). The data is plotted (C) as a histogram, or (D) normalized by the radius to the DMV centre to account for the increase in the screened area. Scale bars, (A) 5 μm, (B) 500 nm.

Next, we specifically investigated the possible involvement in vRNA synthesis of CM and/or DMSs, which always were present in membrane-modification clusters that also contained DMVs. A close inspection of CM showed that these structures were mainly devoid of signal and that the occasional silver grains present primarily appeared in the periphery of CM, therefore likely stemming from surrounding labelled DMVs (Fig 5A and B). Similar observations were made for DMSs (Fig 5C-E): most of them (85%) lacked signal and the rest was close to abundantly labelled DMVs. Furthermore, no signs of DMSs acting as a signal source were apparent in the distribution of signal around them, which resembled that of a random pattern (Fig 5E, compare with Fig 4D).

**Fig. 5.**
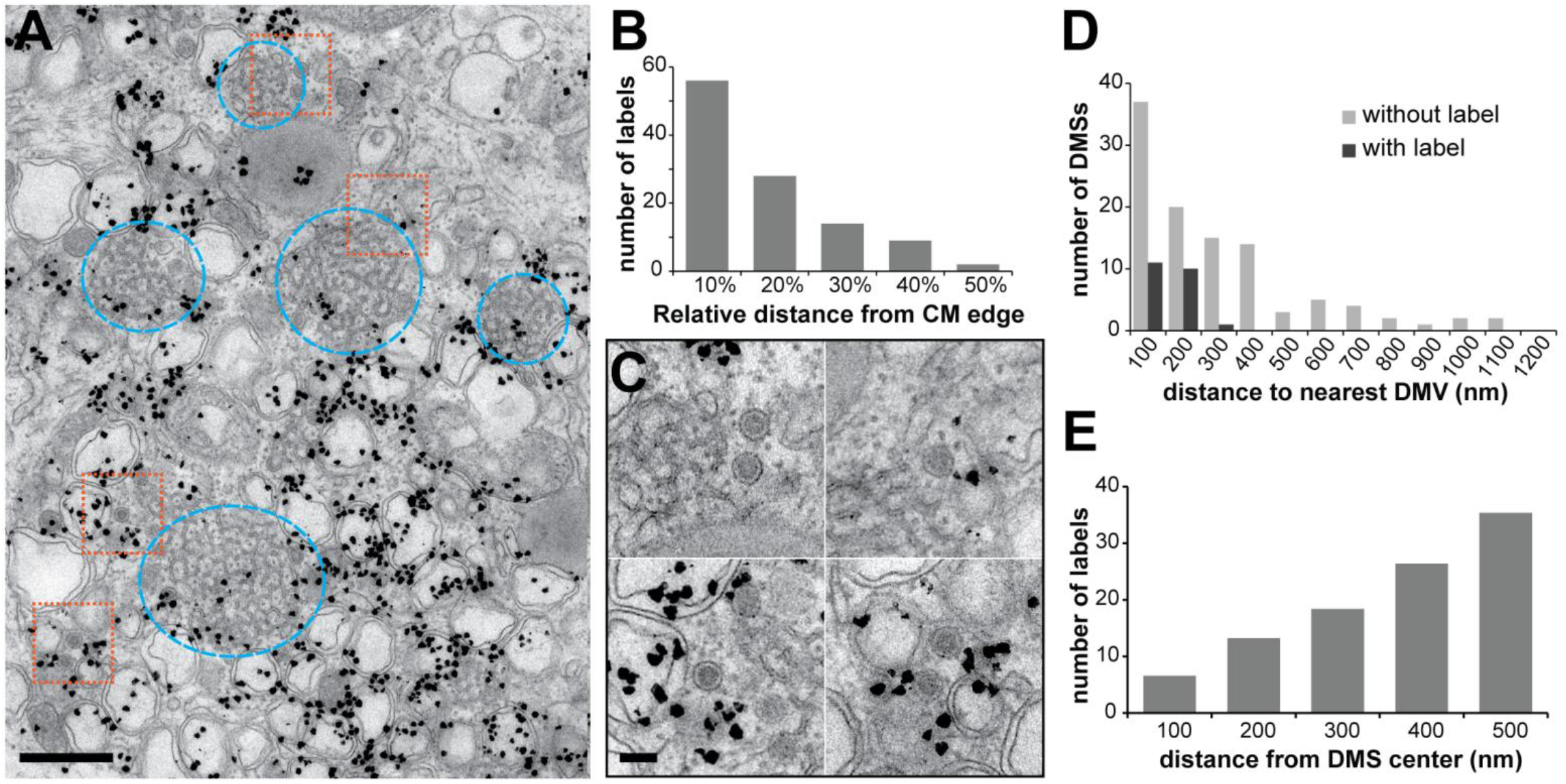
Newly-synthesized vRNA signal does not clearly associate with CM or DMSs. (A) Overview of a cluster of MERS-CoV-induced membrane modifications in Huh7 cells prepared as described in Fig. 4. Some DMSs are boxed in orange while regions with CM are encircled in blue. In comparison with the densely-labelled surrounding DMVs, these regions are relatively devoid of autoradiography signal. (B) The distribution of autoradiography grains on CM was not homogeneous (n_CM_=9), and label was predominantly found close to the boundaries of the CM, as expected if the signal arises from the surrounding DMVs. (C-E) Analysis of the label around/on the DMSs (see Materials and Methods for selection criteria and details). (C) Enlargements of the DMS areas boxed in (A). Most DMSs were devoid of signal and those who contained label were close to labelled DMVs (D) (n_DMS_ = 105). (E) The distribution of signal around DMSs shows an increase in the amount of autoradiography grains with the distance from the DMS centre, as expected from a random distribution (n_DMSs_ = 54). Scale bars, (A) 500 nm, (C) 100 nm.

To explore whether these observations could be extended to distantly-related CoVs, we expanded this type of analysis to the gamma-CoV IBV. IBV-induced DMSs are particularly abundant and a large proportion of them have an open configuration, which contributed to the hypothesis that these open DMSs could be engaged in vRNA synthesis [29, 31]. However, no evidence supporting this hypothesis could be derived from our detailed analysis of the autoradiography signal in IBV-infected cells, which essentially produced the same results as for MERS-CoV (S3 Fig). Taken together, our observations clearly point at coronaviral DMVs as the active site of vRNA synthesis and seem to indicate that neither CM/zippered ER nor DMSs are effectively involved in this process, at least not to a significant level above the detection limits of our method.

### DMSs do not label for viral envelope proteins or dsRNA

If CM and DMSs are not involved in vRNA synthesis, what is the role (if any) of these RO structural elements in coronavirus replication? To investigate this, we analysed the subcellular location of different viral markers in MERS-CoV-infected cells by immunoelectron microscopy (IEM) (Fig 6).

**Fig. 6.**
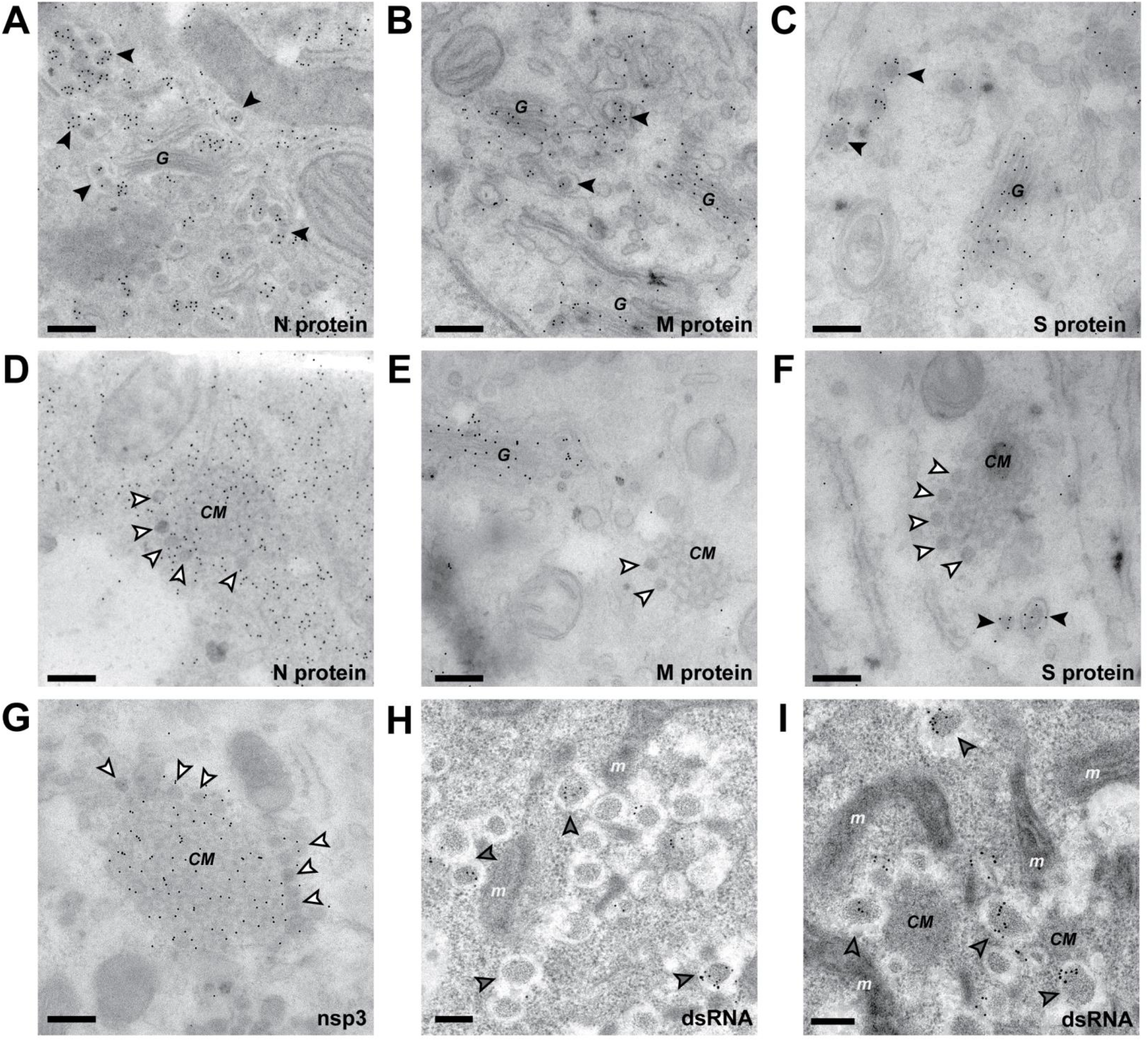
IEM detection of viral markers in MERS-CoV-infected cells. (A-G) Immunogold labeling of thawed cryo-sections of MERS-CoV-infected Huh7 cells (12 hpi) for the detection of the indicated viral proteins. (A-C) Structural proteins were detected on virions (black arrowheads) and, for the M and S proteins, also on Golgi cisterna. While regions containing DMS (white arrowheads) and CM labelled for the N protein (D) and nsp3 (G), the M and S protein were not detected in these areas. (H-I) Immunogold labeling of dsRNA in HPF-FS samples of MERS-CoV-infected Huh7 cells (13 hpi). The label accumulated on DMVs, which could be easily detected in this type of samples (grey arrowheads), while the regions with CM and DMSs, which appeared as dark areas among the DMV clusters, where devoid of dsRNA signal. G, Golgi complex; m, mitochondria. Scale bars, 250 nm.

Coronavirus-induced DMSs, with their electron-dense content and their remarkably regular size, are particularly intriguing structures. Revealing DMSs in IEM samples, however, was challenging and, in our hands, required a modified protocol for the preparation of thawed cryo-sections [43] that, unfortunately, failed to make DMVs apparent (see Materials and Methods). Given the similar size of DMSs and virus particles, we first considered the possibility that the DMSs would represent some kind of non-productive virus assembly event on the CM. While new CoV particles typically assemble in the ERGIC [40–42, 44, 45], virus budding from ER membranes, from which CM originate, can also occur [40, 46, 47], and we regularly observed it in MERS-CoV- and IBV-infected cells (e.g. S1 Fig).To investigate this possibility, we used antibodies against several of the structural proteins, namely, the nucleocapsid protein N, the envelope membrane protein M and the spike protein S. As expected, all of them were detected in newly-formed MERS-CoV particles present in budding vesicles (Fig 6A-C).The M and S proteins also localized to the Golgi complex, aligning with previous observations for other CoVs [20, 44, 48, 49]. The MERS-CoV N protein was found in regions with CM and DMSs, though the distribution of signal was homogenous and DMSs were not particularly densely labelled (Fig 6D). The presence of the N protein in the viral RO has also been shown for MHV [20] and suggested by a number of colocalisation studies [42, 50–53], and may be related to a possible role in vRNA synthesis of this multifunctional protein [54]. Importantly, neither DMSs nor CMs labelled for the M protein, the most abundant viral envelope protein and the presumed orchestrator of virion assembly, or the S protein (Fig 6E and 6F).

Previously, the CM induced by SARS-CoV and MHV were shown by IEM to accumulate viral nsps, while dsRNA signal was primarily found inside the DMVs [15, 20]. Similarly, nsp3 mapped to the CM induced in MERS-CoV infection, but also to the DMSs to a comparable extent (Fig 6G). Our attempts to combine dsRNA antibody labelling with thawed cryo-sections were unsuccessful, which made us resort to HPF-FS samples. In these, however, while DMVs were easily detected, the morphology of CM and DMSs was less clearly defined. Nevertheless, dsRNA signal was clearly associated with DMVs, while the dark membranous regions between DMVs that we interpreted as CM and DMSs clusters appeared devoid of signal (Fig 6H and 6I).

In summary, for the antibodies tested (recognizing N, M, S, nsp3, and dsRNA), the labelling pattern in MERS-CoV-induced DMSs closely resembled that of the CM, from which they seem to derive. The absence of labelling for key proteins in virus assembly, like the M and S proteins, strongly suggest that DMSs do not represent (spurious) virus assembly events.

## Discussion

The comprehensive analysis presented here demonstrates that viruses across different CoV genera induce essentially the same type of membrane structures. After somewhat disparate observations [15, 20, 21, 29, 30, 47], the unifying model that emerges from our study is that of a coronavirus RO comprising three basic types of double-membrane structural elements: (i) DMVs, (ii) CM or zippered ER, which would represent branched or un-branched configurations of paired-ER membranes, and (iii) small DMSs that appear to arise from CM or zippered ER. These structural elements are largely interconnected and connected to the ER, together forming the reticulovesicular network that is typical of the coronaviral RO, including in all likelihood the RO of the 2019-nCoV (SARS-CoV-2), a close relative of SARS-CoV.

Our results clearly establish DMVs as the primary ‒if not only-site of vRNA synthesis within the coronaviral RO. This point has been subject to quite some speculation, due in part to the limited experimental data directly addressing this question. An early study using Br-UTP and IEM to detect newly-synthesized vRNA mapped signal in DMV regions of MHV-infected cells [14], but the poor preservation of IEM samples did not allow the recognition of CM and DMSs, typically present in these regions. Later, a light microscopy study using 5-ethynil uridine labelling and click chemistry suggested that vRNA synthesis could take place, at least partially, in a different location than the DMVs [55]. Alternative interpretations of those results, like migration of vRNA from the DMVs during the relatively long labelling times used (60 min) appear now more plausible. Several other studies further contributed to undermine the idea of DMVs being the sites of vRNA synthesis by showing that higher numbers of DMVs did not necessarily translate into larger amounts of vRNA or provide a competitive advantage [27, 28]. It should be noted, however, that the number of DMVs may not necessarily correlate with the number of active replication complexes that they contain at a given time, something that is also suggested by our observed variations in the level of vRNA synthesis signal among DMVs.

While the finding of open DMSs, similar to the invaginations that many +RNA viruses use as replication sites, made them attractive candidate sites for vRNA synthesis [29, 31], we could not detect vRNA synthesis associated with them, nor with CM or any other subcellular structure using a highly-sensitive technique like autoradiography. Although some level of vRNA synthesis in any of these structures cannot be completely discarded, our results suggest that, if present, this would only be marginal compared to the abundant synthesis clearly associated with DMVs. However, given the prevalence of DMSs and CM across different CoV genera, it is tempting to speculate that these structures must play a role in virus replication. It has been proposed that CM could be a form of cubic membranes [25, 56], which are membrane aggregates resulting from ER protein overexpression [57]. CM, which abundantly label for viral nsps [15, 20] and proliferate late in infection [15, 20, 21], could thus be a by-product of viral protein overexpression. Although DMSs seem to be derived from CM, it is harder to imagine that these highly-regular structures would also lack a specific function. However, their possible role remains elusive. The idea that DMSs could be DMV precursors seems unlikely, considering that no intermediate structures between the two were found, and that DMV formation precedes the appearance of CM, from which DMSs seem to originate. A suggestive possibility was that DMSs represented non-productive virus assembly events, but the lack of DMS labelling for key structural proteins seems to rule out this option. In fact, no differences were apparent in the labelling patterns of CM and DMSs for the viral markers tested. While their distinct morphology likely implies that DMSs contain specific host or viral proteins, these factors, which may give important clues about the DMS role, remain to be identified.

Our results add to studies that, in the last years and after much speculation, have started to provide experimental evidence that the DMVs induced by +RNA viruses are active sites of vRNA synthesis [11, 58–60]. However, it is not clear that DMVs always play the primary role in virus replication that we demonstrate here for CoV. For picornaviruses, for example, virus-induced single membrane structures, which are DMV precursors and also active sites of vRNA synthesis, could well be more relevant as they predominate at the peak of vRNA replication [10, 11, 60]. By clearly pointing to DMVs as the key sites for CoV replication, our results also bring back to centre stage some of the challenges that CoV-induced DMVs pose, which extend to the distantly-related arterivirus family, and probably also to other members of the order *Nidovirales*. In contrast with the DMVs induced by other +RNA viruses [10, 11, 13, 60], the DMVs in nidovirus-infected cells appear to lack membrane openings that would connect their inner compartment with the cytosol to allow import of precursors and export of genomic and subgenomic viral mRNAs [15, 16, 29]. This topological conundrum, however, starts with the assumption that vRNA synthesis takes place inside the DMVs, yet the evidence so far is insufficient to ascertain this point. Shielding of vRNA inside DMVs arguably provides the most straightforward explanation for the observation that intact membranes protected vRNA from nuclease treatment [61]. However, protection of vRNA could also be achieved in the outer DMV membrane through protein complexes that would rely on membranes for assembly/stability.

While a definitive answer to this issue is still missing, our results allow narrowing down the possible scenarios. If the DMVs are closed structures lacking an import/export mechanism, vRNA synthesis on the outer DMV membrane appears as a necessity to provide vRNA for translation and encapsidation. Then, vRNA synthesis inside the DMVs would also be required to explain the observed accumulation of (presumably viral) dsRNA. Although this is not at all an attractive possibility, as vRNA synthesis inside the DMVs and even DMV formation would appear spurious, it cannot be completely discarded at this stage. The most appealing scenario is that in which vRNA synthesis only takes place inside the DMVs. This would provide the compartmentalization of vRNA synthesis that may be most beneficial for viral replication, although it would require the existence of a yet unidentified import/export mechanism. Notice that a transport mechanism would also be needed in the third possible scenario, i.e. if vRNA synthesis occurs only on the outer DMV membrane, to account for the accumulation of dsRNA inside the DMVs that could then perhaps serve to hide excess vRNA from detection by innate immune sensors [62].

Despite the lack of clear openings in the membranes of CoV-induced DMVs, a mechanism allowing exchange of material with the cytosol is conceivable. The openings of DMV may be extremely short-lived, and therefore they may have eluded detection by EM. This may become apparent in the future, if mutant CoV inducing DMVs with slower dynamics are found. An appealing alternative is the existence of molecular pores that may well be undetectable in conventional EM samples. Precedents of molecular complexes bridging double membranes include the large nuclear pore complex, but also small transporters through the mitochondrial or chloroplast membranes. Visualizing such a putative small molecular pore on DMVs is a formidable challenge that would likely require the use of cryo-EM to preserve macromolecular components. Emerging techniques like *in situ* cryotomography, which allows the visualization of structures in their cellular context at macromolecular resolution, may be key to realize this next step and to understand how CoVs exploit the complex architecture of DMVs.

## Materials and Methods

### Cells, viruses and infections

Huh7 cells (kindly provided by Ralf Bartenschlager, Heidelberg University) were grown in Dulbecco’s modified Eagle’s medium (DMEM; Lonza) supplemented with 8% (v/v) fetal calf serum (FCS; Bodinco), 2 mM L-glutamine (PAA Laboratories) and nonessential amino acids (PAA Laboratories). Vero cells (ECACC 84113001) were cultured in Eagle’s minimal essential medium (EMEM; Lonza) with 8% FCS and 2 mM L-glutamine. Vero E6 cells (ATCC CRL-1586) were maintained in DMEM supplemented with 8% FCS. Mouse 17 clone 1 (17Cl1) cells (gift from Stuart Siddell, University of Bristol) were grown in DMEM supplemented with 8% FCS and 8% (v/v) tryptose phosphate broth (Life technologies). Penicillin and streptomycin (90 IU/ml, PAA Laboratories) were added to all media.

The coronaviruses used in this study include MERS-CoV (strain EMC/2012, [32, 33], SARS-CoV (strain Frankfurt-1, [63]), MHV (ATCC VR-674), human coronavirus strain 229E (HCoV-229E, [64]), and IBV (strain Beau-R, [65]). All the infection experiments were carried out at 37°C, except for HCoV-229E infections which were performed at 33°C. Cells were infected at high multiplicity of infection (MOI 5-10), with the exception of IBV (MOI<1). In this case, the time post-infection chosen (~16 hpi) allowed for two cycles of IBV replication to maximize the number of infected cells. For MHV infections, 1 μM HR2 peptide was added to the cell medium to prevent syncytia formation [66]. Control mock-infected cells were included in all the experiments. Infections were routinely assessed by immunofluorescence assays on parallel samples, essentially processed as previously described [51]. All work with live SARS-CoV and MERS-CoV was performed inside biosafety cabinets in the biosafety level 3 facility at Leiden University Medical Center.

### Antibodies

For IEM of MERS-CoV-infected cells, the antibodies used included a previously described rabbit antiserum that recognizes SARS-CoV nsp3 protein and cross reacts with MERS-CoV nsp3 [21, 67], a polyclonal rabbit antibody generated against full-length MERS-CoV N protein (Sino Biological), and a mouse monoclonal antibody (J2) specific for dsRNA [68], purchased from Scicons. A MERS-CoV M-specific rabbit antiserum was ordered from Genscript and produced using as antigen a synthetic peptide representing the C-terminal 24 residues of the protein (CRYKAGNYRSPPITADIELALLRA). The specificity of the antiserum was verified by Western blot analysis and immunofluorescence microscopy using samples from MERS-CoV-infected Vero or Huh7 cells, as described previously [21], whilst using pre-immune serum and mock-infected cell lysates as negative controls. The human monoclonal antibody used against MERS-CoV spike protein was kindly provided by Dr. Berend Jan Bosch (Utrecht University) and has been described elsewhere as 1.6f9 [69].

### Metabolic labelling and label incorporation measurements

To label newly-synthesized RNA, CoV-infected cells and control mock-infected cells were incubated for different periods of time with tritiated uridine ([5-^3^H]uridine, 1 mCi/ml, Perkin Elmer), which was mixed in a 1:1 ratio with double-concentrated medium. Cellular transcription was blocked by providing the cells with 10 μg/ml actinomycin D both during labelling and in a pre-incubation step of 1-2 h, depending on the specific set of samples. Directly after labelling, the cells were extensively washed with phosphate-buffered saline (PBS) and immediately fixed for EM autoradiography. A parallel set of samples was included in every experiment for label incorporation measurements. These cells were lysed using TriPure Isolation Reagend (Roche) and the RNA isolated following the manufacturer’s instructions. The incorporation radioactive label into RNA was then measured using scintillation counting.

### Sample preparation for electron microscopy

For ultrastructural analysis, autoradiography and tomography, EM samples of CoV-infected cells were prepared by chemical fixation. Chemical fixation was chosen over the alternative of high-pressure freezing (HPF) and freeze-substitution (FS) to increase the yield of cells per EM grid and thus facilitate the autoradiography quantitative analysis, as it was observed that infected cells were easily washed away and lost during FS. At the desired times post infection, the cells were fixed for 30 min with 1.5% (vol/vol) glutaraldehyde (GA) in 0.1 M cacodylate buffer (pH 7.4). Cells infected with SARS- or MERS-CoV were further maintained in the fixative overnight. After fixation, the samples were washed with 0.1 M cacodylate buffer either once or, in samples destined to autoradiography, 3 times (5 min each) to favour the elimination of unincorporated [5-^3^H]uridine. Next, the samples were treated with 1% (wt/vol) OsO_4_ at 4°C for 1 hour, washed with 0.1 M cacodylate buffer and Milli-Q water, and stained at room temperature for 1 hour with 1% (wt/vol) low-molecular weight tannic acid (Electron Microscopy Science) in 01. M cacodylate buffer (SARS-CoV autoradiography experiments), or with 1% (wt/vol) uranyl acetate in Milli-Q (rest of the samples). Following a new washing step with Milli-Q water, the samples were dehydrated in increasing concentrations of ethanol (70%, 80%, 90%, 100%), embedded in epoxy resin (LX-112, Ladd Research), and polymerized at 60°C. Sections were collected on mesh-100 copper EM grids covered with a carbon-coated Pioloform layer, and post-stained with 7% uranyl acetate and Reynold’s lead citrate.

#### EM autoradiography samples

To facilitate the different steps in the preparation of autoradiography samples, EM grids with ultrathin cell sections (50 nm thick) were first attached to glass slides by gently pressing the grid edge against a line of double-sided sticky tape applied along the length of the slide. Each glass slide included a grid for every condition tested within a given experiment, to be later developed simultaneously. A thin layer of carbon was then evaporated onto the grids to prevent direct contact between the stained section and the photographic emulsion that could lead to undesired chemical changes in the emulsion. Samples were transferred to a dark room, and a thin layer of nuclear emulsion ILFORD L4 was placed on top of the grids with the help of a wire loop [36]. Samples were maintained in the dark in a cold room for several weeks until development, which was performed as described in [70]. The progress in the exposure of the nuclear emulsion to radioactive disintegrations was evaluated regularly by EM until the number of autoradiography grains was sufficient for analysis (~6 weeks for SARS-CoV-infected cells, ~21 weeks for MERS-CoV-infected cells, and ~12-13 weeks for IBV-infected cells).

#### IEM

Several types of MERS-CoV-infected cell samples were prepared for IEM. The labelling of viral proteins was performed on thawed cryo-sections, which are optimal for epitope preservation. To this end, at 12 hpi, infected and mock-infected cells were first chemically fixed for 1 hour at room temperature with 3% (wt/vol) paraformaldehyde (PFA) and 0.25% (vol/vol) GA in 0.1 M PHEM buffer (60 mM PIPES, 25 mM HEPES, 10 mM EGTA, 2 mM MgCl_2_, ph 6.9), briefly washed with 0.1 M PHEM buffer and stored at 4°C in 3% (wt/vol) PFA in 0.1 M PHEM buffer until transfer from the biosafety level 3 facility for further processing. To prepare EM samples, the cells were first pelleted and embedded in 12% (wt/vol) gelatin. Cubes of approximately 1 mm^3^ in size were cut from these pellets, infiltrated with 2.3 M sucrose for cryo-protection, **s**nap-frozen in liquid nitrogen, and sectioned by cryo-ultramicrotomy. Thawed cryo-sections (70 nm thick) deposited on EM grids were incubated first with the corresponding primary antibody and then with protein A coupled to colloidal 10-nm gold particles. However, virus-induced membrane modifications were not discernible in these samples, and clear signs of membrane extraction were present. To tackle this issue, we used a previously described modification of the protocol that includes sequential post-staining steps with 1% 1% (wt/vol) OsO4, 2% (wt/vol) uranyl acetate and Reynold’s lead citrate after immunogold labelling and prior to the final embedding in a thin layer of 1.8% methyl cellulose [43]. Both CM and DMSs were clearly recognizable in these samples, while DMVs were still not apparent and may have been extracted, as empty areas were often found in the vicinity of CM regions.

The detection of dsRNA required the preparation of HPF-FS samples. For biosafety considerations, Huh7 cells grown on sapphire disks and infected with MERS-CoV, were first fixed overnight at 13 hpi with 3% (wt/vol) paraformaldehyde and 0.25% (vol/vol) GA in 0.1 M PHEM buffer. Then, the samples were frozen with a Leica EM PACT2, after which they were freeze-substituted in a Leica AFS2 system with 0.1% (wt/vol) uranyl acetate as previously described [71], with the only modification that acetone was replaced by ethanol from the last washing step before Lowycril infiltration onwards. Cell sections (75 nm thick) were incubated with the primary mouse antibody, then with a bridging rabbit anti-mouse-IgG antibody (Dako Cytomation), and finally with protein A coupled to 15-nm gold particles. After immunolabelling, samples were additionally stained with 7% uranyl acetate and Reynold’s lead citrate.

#### Electron Microscopy Imaging

Individual 2D-EM images were acquired in a Tecnai12 BioTwin or a Twin electron microscope, equipped with an Eagle 4k slow-scan change-couple device (CCD) camera (Thermo Fisher Scientific (formerly FEI)) or a OneView 4k high-frame rate CMOS camera (Gatan), respectively. Mosaic EM images of large grid areas were generated for the quantitative analysis of autoradiography samples, using overlapping automatically-collected images (pixel size 2 nm, Tecnai 12 BioTwin) that were subsequently combined in a composite image as described in [38].

#### Electron Tomography

Prior to the post-staining step, semi-thin sections (150 nm thick) of CoV-infected cells were incubated with protein A coupled to 10-nm colloidal gold particles that served later as fiducial markers for alignment. Dual-axis tilt series were collected in a Tecnai12 BioTwin microscope using Xplore 3D acquisition software (Thermo Fisher Scientific), covering each 120-130° around the specimen with an angular sampling of 1° and a pixel size of 1.2 nm. The alignment of the tilt series and tomogram reconstruction by weighted back-projection was carried out in IMOD [72]. The diameters of DMVs, DMSs and virions were measured at their equator. DMV profiles, often only roughly circular, were measured over their longest and shortest axes, and the diameter was estimated as the geometric mean of the two values. For visualization purposes, the tomograms were first mildly denoised and then processed in Amira 6.0.1 (Thermo Fisher Scientific) using a semi-automatic segmentation protocol as previously described [10].

#### Quantitative analysis of autoradiography samples

Large mosaic EM maps containing dozens of cell profiles were used for the quantitative analysis of the newly-synthesized RNA autoradiography signal (see S1 Table). For each CoV, different conditions (infected and mock-infected cells, plus different labelling times) were compared using only samples developed after the same period of time. The analysis of the signal in different subcellular regions was carried out using home-built software. Areas of 4 μm^2^ were randomly selected from the mosaic EM maps and the autoradiography grains present in those areas were manually assigned to the underlying cellular structures. The abundance of the different types of subcellular structures was estimated through virtual points in a 5×5 lattice superimposed to each selected area, which were also assigned to the different subcellular classes. Regularly along the process, the annotated data per condition was split into two random groups and the Kendall and Spearman coefficients, which measure the concordance between two data sets [73], were calculated. New random regions were added until the average Kendall and Spearman coefficients resulting from 10 random splits were higher than 0.8 and 0.9, respectively (maximum value, 1). Labelling densities and relative labelling indexes (RLI) were then calculated from the annotated points [39].

For the analysis of the association of vRNA synthesis with each of the different ROs motifs, the specific DMVs, DMSs and CM included in the analysis were carefully selected. Only individual DMVs that were at least one micron away from any other virus-induced membrane modification were included in the analysis. For every grain present in an area of 750 nm radius around each DMV, the distance to the DMV centre was measured. In the case of DMSs, which were always part of clusters of virus-induced membrane structures, only DMSs in the periphery of these clusters were selected. The quantified signal was limited to sub-areas devoid of other RO motifs, which were defined by circular arcs (typically 30° to 100°, radius 500 nm) opposite to the RO clusters. CMs are irregular structures that appear partially or totally surrounded by DMVs. Only large CM (> 0.6 μm across) were selected in order to make more apparent (if present) any decay of the autoradiography signal as the distance to the surrounding DMVs increased. For each autoradiography grain, both the distance to the closest CM boundary (*d_1_*) and the distance to the opposite CM edge (*d_2_*) were measured. The relative distance to the CM edge was then calculated as *d_1_/(d_1_+d_2_)* and expressed in percentages. All the measurements in different DMVs, DMSs and CM were made using Aperio Imagescope software (Leica) and pooled together into three single data sets.

## Acknowledgments

This work was supported by the Netherlands Organization for Scientific Research (NWO-MEERVOUD-836.10.003 to M.B.). We are grateful to Yvonne van der Meer, Charlotte Melia, and Barbara van der Hoeven for their assistance.

## Supporting Information

### S1 Appendix: EM autoradiography

Autoradiography is a classic technique that allows the EM visualization of a radioactive marker, usually targeting a certain process, and thus reveals the subcellular localization of that process [1, 2]. Tritiated uridine, for example, can be used to locate active RNA synthesis [3–5], as shown also in this study. A clear advantage over the use of alternatives for metabolic labelling of newly-synthesized RNA (e.g. Br-uridine, Br-UTP, 5-ethynil uridine) is that the radioactive precursor is chemically identical to the natural substrate.

After labelling, the samples are immediately fixed and processed for EM. The location of the radioactive marker can then be made apparent by applying a highly-sensitive photographic emulsion (a nuclear emulsion) on top of the cell sections and exposing it for several weeks to months. The beta particles that are emitted as a result of tritium disintegrations generate electrons that get trapped in the silver halide emulsion and create a “latent image”. When the emulsion is developed, these negative charges promote the reduction to metallic silver, generating electron-dense grains that are visible by EM. In principle, given enough time to accumulate enough radioactive disintegrations, even low levels of the radioactive marker could be detected. In practice, other factors (e.g. background radiation, emulsion aging) set some limits to autoradiography, which is nonetheless a very sensitive technique.

The resolution of EM autoradiography is limited by the fact that radioactive disintegrations generate beta particles that are emitted in random directions. Importantly, the probability of giving rise to signal degreases with the distance from the radioactive source; however, some beta particles may travel up to a few hundred nanometers before striking the photographic emulsion [2]. Therefore, it is important to keep in mind that the silver grains may not directly overlay the structure containing the radioactive source. Quantitative analyses of the signal that take this factor into account, like those presented in this study, become indispensable to maximize the information that autoradiography can provide.

## Supporting Figures

**S1 Fig.**
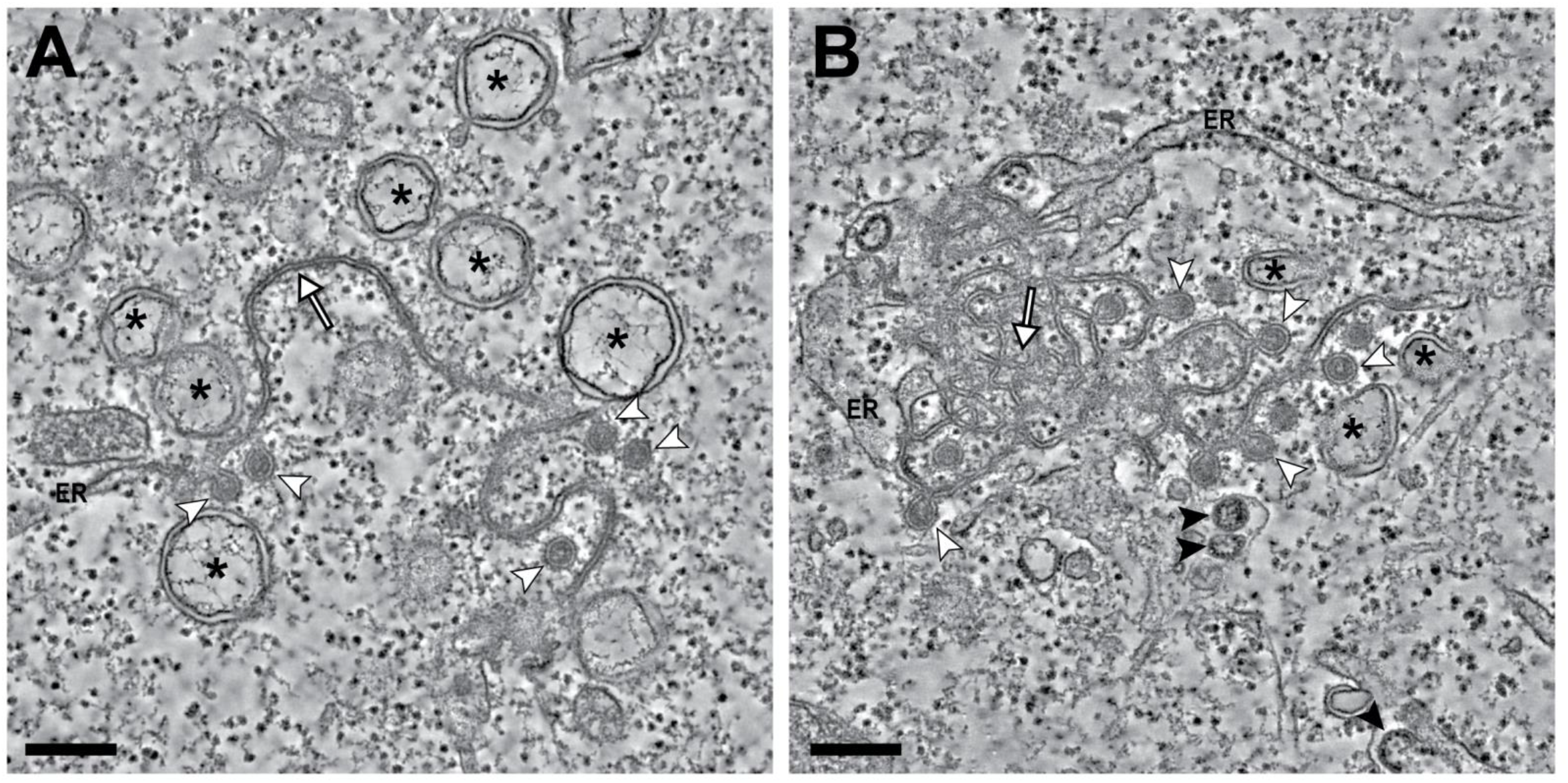
Membrane structures induced by gamma-CoV infections. Tomography of Vero-E6 cells infected with IBV, fixed at 18 hpi and processed for EM following the same protocol as for MER-CoV-infected cells (Fig 1). Tomographic slices through two regions containing IBV-induced membrane modifications. These include DMVs (asterisks), DMSs (white arrowheads) and zippered ER (white arrows). Most zippered ER consists of long stretches of ER-derived paired membranes (A), though branching zippered ER, closer to the CM described for beta-CoV, was also present (B) Virus particles (black arrowheads) budding into the ER membranes were often observed. Scale bars, 250 nm.

**S2 Fig.**
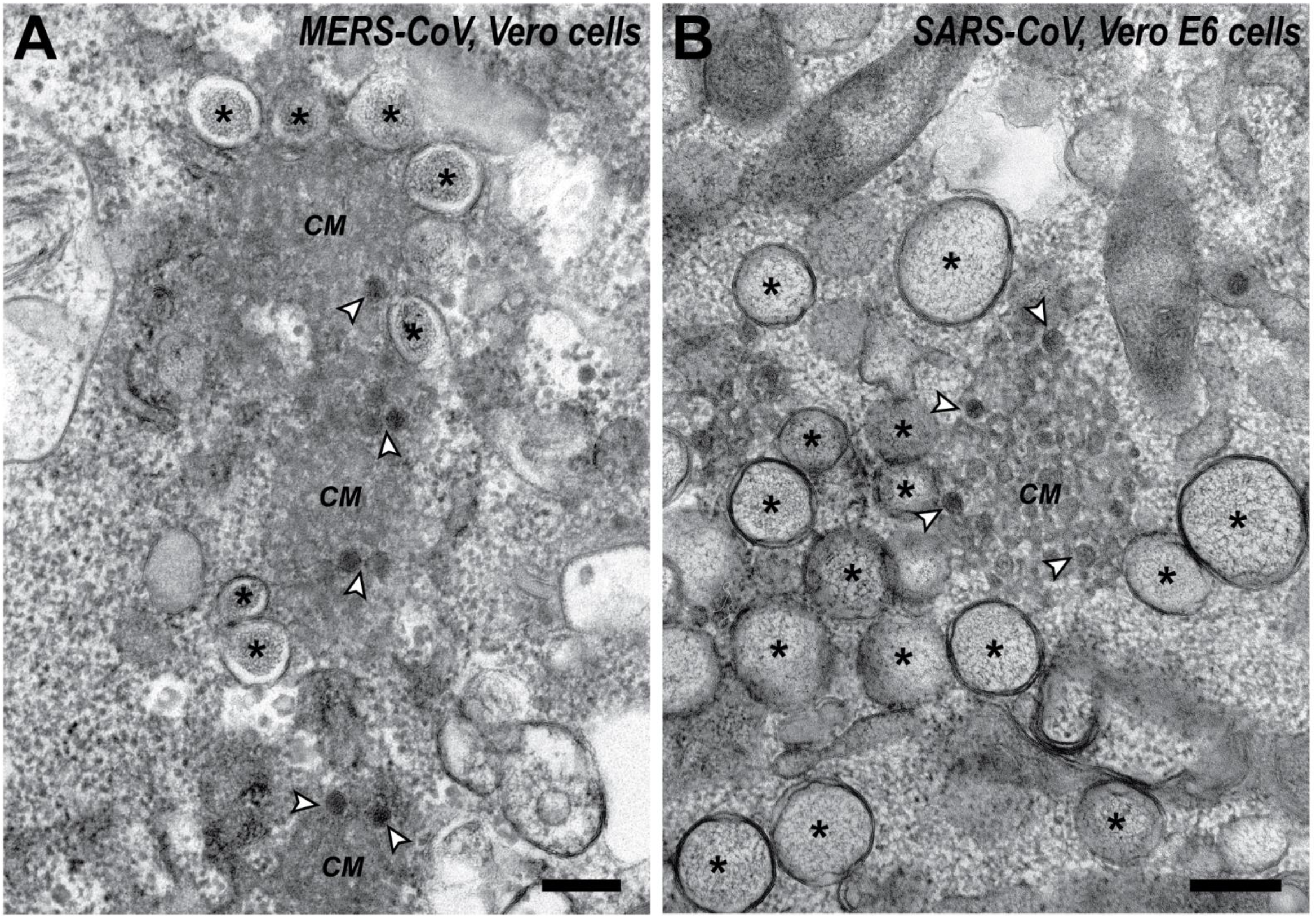
Detection of DMSs in cryo-fixed and FS samples of CoV-infected cells. Analysis of previously described samples of CoV-infected cells, prepared for EM either by HPF(A) or cryo-plunging (B). A targeted search revealed the presence of DMSs (white arrowheads) in close association with CM. In comparison with the chemically fixed samples used in this study, the superior ultrastructural preservation of cryo-fixation results in less distorted membranes, but also in a denser cytoplasm and darker CM that makes DMS less apparent. (A) Example from a MERS-CoV-infected Huh7 cell (16 hpi) in a sample used in [6]. (B) Region in a SARS-CoV-infected cell (8 hpi), adapted from [7]. Scale bars, 250 nm.

**S3. Fig.**
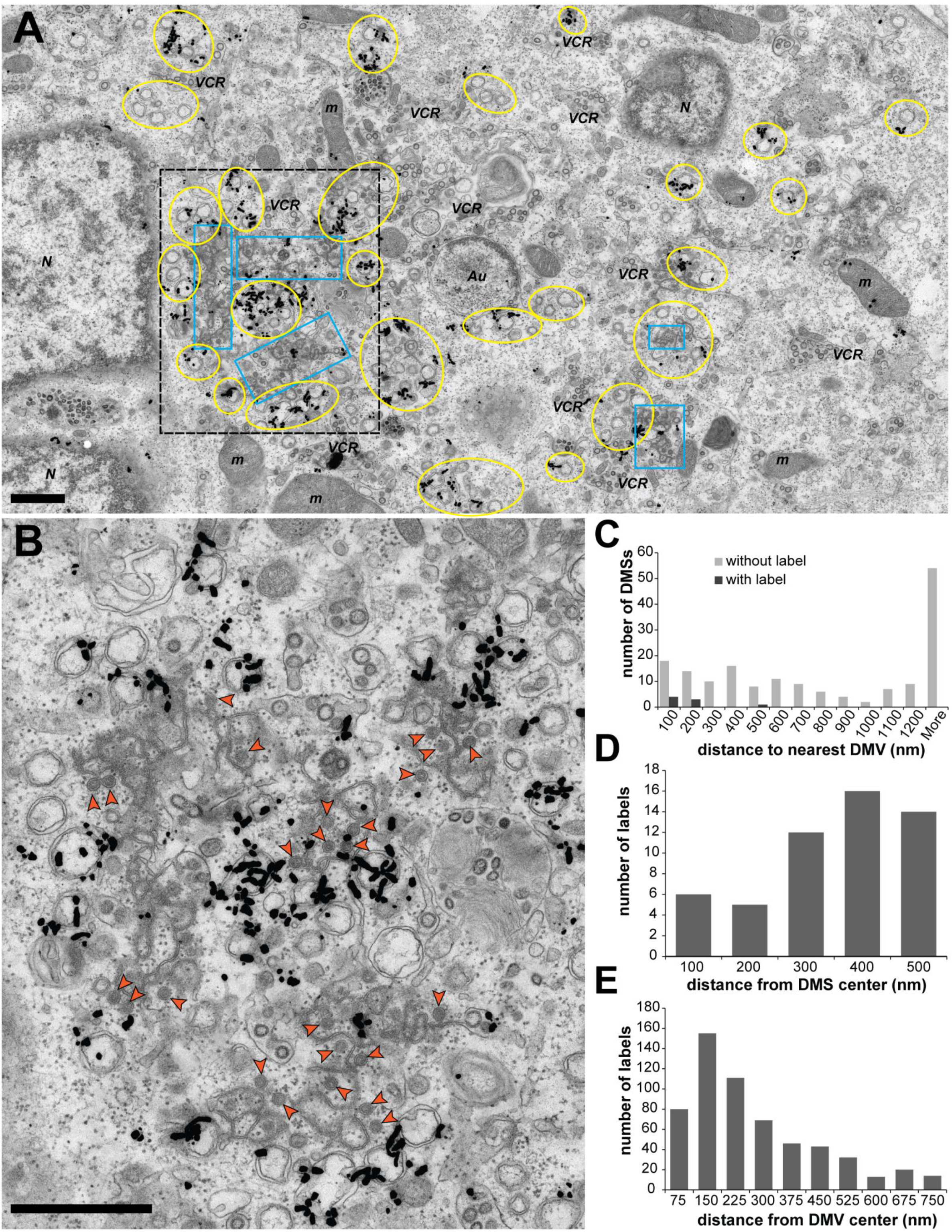
Metabolic labelling of newly-synthesized vRNA in IBV-infected cells and analysis of the autoradiography signal. Vero cells infected with IBV were pre-treated with actinomycin D for 1 hour, then labelled for 30 or 60 min with tritiated uridine, immediately fixed at 16 and 17 hpi, respectively, and processed for autoradiography EM. (A) Overview of an IBV-infected Vero cell (60 min labelling). The areas containing DMVs and zippered ER are outlined in yellow and blue, respectively, and other subcellular structures annotated (N, nucleus; m, mitochondria; Au, autophagosome; VCR, virion-containing regions). The autoradiography signal accumulates in areas of virus-induced membrane modifications that often only contain DMVs, in alignment with DMVs having an active role in vRNA synthesis. (B) Close-up of the area boxed in black in (A), which contains DMVs, zippered ER and DMSs (orange arrowheads). The contrast between the densely labelled DMVs and the zippered ER and DMSs largely lacking signal is apparent and suggests that the autoradiography grains sometimes present on the latter structures arose from radioactive disintegrations in the surrounding active DMVs. (C) In agreement with this possibility, most of the DMSs (96%) were devoid of signal, and most of those that contained label where close to an active DMV (n_DMS_ = 178). (D) Furthermore, the distribution of autoradiography grains around DMSs resembled that of a random distribution, where the number of grains increase with the distance (n_DMS_ =106). (E) In contrast, a similar analysis of the signal around the DMVs proved that these structures are associated with vRNA synthesis, as the signal reaches maximum values in the proximity of the DMVs (n_DMVs_ = 51). ((C, D) See Materials and Methods for the selection criteria and details). Scale bars, 1 μm.

### S1 Video

#### Electron tomography of the membrane structures induced in MERS-CoV infection

Animation illustrating the tomography reconstruction and model presented in Fig 1B. The video first shows the tomographic slices (1.2 nm thick) through the reconstructed volume, and then surface-rendered models of the different structures segmented from the tomogram: DMSs (orange), CM (blue) and DMVs (yellow and lilac, outer and inner membranes), ER (green), and a vesicle (silver) containing virions (pink). The movie highlights the DMS association with CM, which in turn connect to ER membranes, and these to DMVs.

**S1 Table.** Data sets and sampling for the quantifications of the autoradiography signal presented in Fig 3C and 3D.

|  | Labelling | Data set | Annotations<sup>a</sup> |  |  |
| --- | --- | --- | --- | --- | --- |
|  |  | Cell profiles | Regions | Sampled cellular area (μm<sup>2</sup>) | Labels |
| SARS-CoV-infected Vero-E6 cells | 20 min | 235 | 351 | 1404 | 1367 |
| SARS-CoV-infected Vero-E6 cells | 60 min | 293 | 212 | 848 | 3015 |
| Mock-infected Vero-E6 cells | 60 min | 104 | 436 | 1744 | 607 |
| MERS-CoV-infected Huh7 cells | 30 min | 238 | 210 | 840 | 2324 |
| MERS-CoV-infected Huh7 cells | 60 min | 123 | 211 | 844 | 2233 |
| Mock-infected Huh7 cells | 60 min | 81 | 205 | 820 | 2219 |

<sup>a</sup> numbers calculated excluding those subareas in the randomly selected regions that were outside of the cell

## References

1. Romero-Brey I, Bartenschlager R. Membranous replication factories induced by plus-strand RNA viruses. Viruses. 2014;6(7):2826–57. doi: 10.3390/v6072826. PubMed PMID: 25054883; PubMed Central PMCID: PMC4113795.

2. Harak C, Lohmann V. Ultrastructure of the replication sites of positive-strand RNA viruses. Virology. 2015;479-480:418–33. doi: 10.1016/j.virol.2015.02.029. PubMed PMID: 25746936.

3. Nagy PD, Strating JR, van Kuppeveld FJ. Building Viral Replication Organelles: Close Encounters of the Membrane Types. PLoS Pathog. 2016;12(10):e1005912. Epub 2016/10/28. doi: 10.1371/journal.ppat.1005912. PubMed PMID: 27788266; PubMed Central PMCID: PMCPMC5082816.

4. Scutigliani EM, Kikkert M. Interaction of the innate immune system with positive-strand RNA virus replication organelles. Cytokine & growth factor reviews. 2017;37:17–27. doi: 10.1016/j.cytogfr.2017.05.007. PubMed PMID: 28709747.

5. Schwartz M, Chen J, Janda M, Sullivan M, den Boon J, Ahlquist P. A positive-strand RNA virus replication complex parallels form and function of retrovirus capsids. Molecular cell. 2002;9(3):505–14. PubMed PMID: 11931759.

6. Kopek BG, Perkins G, Miller DJ, Ellisman MH, Ahlquist P. Three-dimensional analysis of a viral RNA replication complex reveals a virus-induced mini-organelle. PLoS biology. 2007;5(9):e220. doi: 10.1371/journal.pbio.0050220. PubMed PMID: 17696647; PubMed Central PMCID: PMC1945040.

7. Welsch S, Miller S, Romero-Brey I, Merz A, Bleck CK, Walther P, et al. Composition and three-dimensional architecture of the dengue virus replication and assembly sites. Cell host & microbe. 2009;5(4):365–75. doi: 10.1016/j.chom.2009.03.007. PubMed PMID: 19380115.

8. Kallio K, Hellstrom K, Balistreri G, Spuul P, Jokitalo E, Ahola T. Template RNA length determines the size of replication complex spherules for Semliki Forest virus. J Virol. 2013;87(16):9125–34. doi: 10.1128/JVI.00660-13. PubMed PMID: 23760239; PubMed Central PMCID: PMC3754052.

9. Fernandez de Castro I, Fernandez JJ, Barajas D, Nagy PD, Risco C. Three-dimensional imaging of the intracellular assembly of a functional viral RNA replicase complex. Journal of cell science. 2017;130(1):260–8. doi: 10.1242/jcs.181586. PubMed PMID: 27026525.

10. Limpens RW, van der Schaar HM, Kumar D, Koster AJ, Snijder EJ, van Kuppeveld FJ, et al. The transformation of enterovirus replication structures: a three-dimensional study of single- and double-membrane compartments. mBio. 2011;2(5). doi: 10.1128/mBio.00166- PubMed PMID: 21972238; PubMed Central PMCID: PMC3187575.

11. Belov GA, Nair V, Hansen BT, Hoyt FH, Fischer ER, Ehrenfeld E. Complex dynamic development of poliovirus membranous replication complexes. J Virol. 2012;86(1):302–12. doi: 10.1128/JVI.05937-11. PubMed PMID: 22072780; PubMed Central PMCID: PMC3255921.

12. Doerflinger SY, Cortese M, Romero-Brey I, Menne Z, Tubiana T, Schenk C, et al. Membrane alterations induced by nonstructural proteins of human norovirus. PLoS Pathog. 2017;13(10):e1006705. doi: 10.1371/journal.ppat.1006705. PubMed PMID: 29077760; PubMed Central PMCID: PMC5678787.

13. Romero-Brey I, Merz A, Chiramel A, Lee JY, Chlanda P, Haselman U, et al. Three-dimensional architecture and biogenesis of membrane structures associated with hepatitis C virus replication. PLoS Pathog. 2012;8(12):e1003056. doi: 10.1371/journal.ppat.1003056. PubMed PMID: 23236278; PubMed Central PMCID: PMC3516559.

14. Gosert R, Kanjanahaluethai A, Egger D, Bienz K, Baker SC. RNA replication of mouse hepatitis virus takes place at double-membrane vesicles. J Virol. 2002;76(8):3697–708. PubMed PMID: 11907209; PubMed Central PMCID: PMC136101.

15. Knoops K, Kikkert M, Worm SH, Zevenhoven-Dobbe JC, van der Meer Y, Koster AJ, et al. SARS-coronavirus replication is supported by a reticulovesicular network of modified endoplasmic reticulum. PLoS biology. 2008;6(9):e226. doi: 10.1371/journal.pbio.0060226. PubMed PMID: 18798692; PubMed Central PMCID: PMC2535663.

16. Knoops K, Barcena M, Limpens RW, Koster AJ, Mommaas AM, Snijder EJ. Ultrastructural characterization of arterivirus replication structures: reshaping the endoplasmic reticulum to accommodate viral RNA synthesis. J Virol. 2012;86(5):2474–87. doi: 10.1128/JVI.06677-11. PubMed PMID: 22190716; PubMed Central PMCID: PMC3302280.

17. Maier HJ, Neuman BW, Bickerton E, Keep SM, Alrashedi H, Hall R, et al. Extensive coronavirus-induced membrane rearrangements are not a determinant of pathogenicity. Scientific reports. 2016;6:27126. doi: 10.1038/srep27126. PubMed PMID: 27255716; PubMed Central PMCID: PMC4891661.

18. Zhang W, Chen K, Zhang X, Guo C, Chen Y, Liu X. An integrated analysis of membrane remodeling during porcine reproductive and respiratory syndrome virus replication and assembly. PloS one. 2018;13(7):e0200919. doi: 10.1371/journal.pone.0200919. PubMed PMID: 30040832; PubMed Central PMCID: PMC6057628.

19. Weber F, Wagner V, Rasmussen SB, Hartmann R, Paludan SR. Double-stranded RNA is produced by positive-strand RNA viruses and DNA viruses but not in detectable amounts by negative-strand RNA viruses. J Virol. 2006;80(10):5059–64. doi: 10.1128/JVI.80.10.5059-5064.2006. PubMed PMID: 16641297; PubMed Central PMCID: PMC1472073.

20. Ulasli M, Verheije MH, de Haan CA, Reggiori F. Qualitative and quantitative ultrastructural analysis of the membrane rearrangements induced by coronavirus. Cellular microbiology. 2010;12(6):844–61. doi: 10.1111/j.1462-5822.2010.01437.x. PubMed PMID: 20088951.

21. de Wilde AH, Raj VS, Oudshoorn D, Bestebroer TM, van Nieuwkoop S, Limpens RW, et al. MERS-coronavirus replication induces severe in vitro cytopathology and is strongly inhibited by cyclosporin A or interferon-alpha treatment. J Gen Virol. 2013;94(Pt 8):1749–60. Epub 2013/04/27. doi: 10.1099/vir.0.052910-0. PubMed PMID: 23620378; PubMed Central PMCID: PMCPMC3749523.

22. Angelini MM, Akhlaghpour M, Neuman BW, Buchmeier MJ. Severe acute respiratory syndrome coronavirus nonstructural proteins 3, 4, and 6 induce double-membrane vesicles. mBio. 2013;4(4). doi: 10.1128/mBio.00524-13. PubMed PMID: 23943763; PubMed Central PMCID: PMC3747587.

23. Hagemeijer MC, Monastyrska I, Griffith J, van der Sluijs P, Voortman J, van Bergen en Henegouwen PM, et al. Membrane rearrangements mediated by coronavirus nonstructural proteins 3 and 4. Virology. 2014;458-459:125–35. doi: 10.1016/j.virol.2014.04.027. PubMed PMID: 24928045.

24. Beachboard DC, Anderson-Daniels JM, Denison MR. Mutations across murine hepatitis virus nsp4 alter virus fitness and membrane modifications. J Virol. 2015;89(4):2080–9. doi: 10.1128/JVI.02776-14. PubMed PMID: 25473044; PubMed Central PMCID: PMC4338892.

25. Oudshoorn D, Rijs K, Limpens R, Groen K, Koster AJ, Snijder EJ, et al. Expression and Cleavage of Middle East Respiratory Syndrome Coronavirus nsp3-4 Polyprotein Induce the Formation of Double-Membrane Vesicles That Mimic Those Associated with Coronaviral RNA Replication. mBio. 2017;8(6). doi: 10.1128/mBio.01658-17. PubMed PMID: 29162711; PubMed Central PMCID: PMC5698553.

26. Doyle N, Neuman BW, Simpson J, Hawes PC, Mantell J, Verkade P, et al. Infectious Bronchitis Virus Nonstructural Protein 4 Alone Induces Membrane Pairing. Viruses. 2018;10(9). doi: 10.3390/v10090477. PubMed PMID: 30200673; PubMed Central PMCID: PMC6163833.

27. Al-Mulla HM, Turrell L, Smith NM, Payne L, Baliji S, Zust R, et al. Competitive fitness in coronaviruses is not correlated with size or number of double-membrane vesicles under reduced-temperature growth conditions. mBio. 2014;5(2):e01107-13. doi: 10.1128/mBio.01107-13. PubMed PMID: 24692638; PubMed Central PMCID: PMC3977362.

28. Lundin A, Dijkman R, Bergstrom T, Kann N, Adamiak B, Hannoun C, et al. Targeting membrane-bound viral RNA synthesis reveals potent inhibition of diverse coronaviruses including the middle East respiratory syndrome virus. PLoS Pathog. 2014;10(5):e1004166. Epub 2014/05/31. doi: 10.1371/journal.ppat.1004166. PubMed PMID: 24874215; PubMed Central PMCID: PMCPMC4038610.

29. Maier HJ, Hawes PC, Cottam EM, Mantell J, Verkade P, Monaghan P, et al. Infectious bronchitis virus generates spherules from zippered endoplasmic reticulum membranes. mBio. 2013;4(5):e00801–13. doi: 10.1128/mBio.00801-13. PubMed PMID: 24149513; PubMed Central PMCID: PMC3812713.

30. Doyle N, Hawes PC, Simpson J, Adams LH, Maier HJ. The Porcine Deltacoronavirus Replication Organelle Comprises Double-Membrane Vesicles and Zippered Endoplasmic Reticulum with Double-Membrane Spherules. Viruses. 2019;11(11). Epub 2019/11/07. doi: 10.3390/v11111030. PubMed PMID: 31694296.

31. Neuman BW, Angelini MM, Buchmeier MJ. Does form meet function in the coronavirus replicative organelle? Trends in microbiology. 2014;22(11):642–7. doi: 10.1016/j.tim.2014.06.003. PubMed PMID: 25037114.

32. Zaki AM, van Boheemen S, Bestebroer TM, Osterhaus AD, Fouchier RA. Isolation of a novel coronavirus from a man with pneumonia in Saudi Arabia. The New England journal of medicine. 2012;367(19):1814–20. doi: 10.1056/NEJMoa1211721. PubMed PMID: 23075143.

33. van Boheemen S, de Graaf M, Lauber C, Bestebroer TM, Raj VS, Zaki AM, et al. Genomic characterization of a newly discovered coronavirus associated with acute respiratory distress syndrome in humans. mBio. 2012;3(6). doi: 10.1128/mBio.00473-12. PubMed PMID: 23170002; PubMed Central PMCID: PMC3509437.

34. Zhu N, Zhang D, Wang W, Li X, Yang B, Song J, et al. A Novel Coronavirus from Patients with Pneumonia in China, 2019. The New England journal of medicine. 2020. Epub 2020/01/25. doi: 10.1056/NEJMoa2001017. PubMed PMID: 31978945.

35. Lu R, Zhao X, Li J, Niu P, Yang B, Wu H, et al. Genomic characterisation and epidemiology of 2019 novel coronavirus: implications for virus origins and receptor binding. Lancet. 2020. Epub 2020/02/03. doi: 10.1016/S0140-6736(20)30251-8. PubMed PMID: 32007145.

36. Bienz KA. Techniques and applications of autoradiography in the light and electron microscope. Microsc Acta. 1977;79(1):1–22. Epub 1977/01/01. PubMed PMID: 65723.

37. Bozzola JJ, Russell LD. Autoradiography & Radioautography. Electron Microscopy: Principles and Techniques for Biologists. Sudbury, MA.: Jones and Bartlett Publishers, Inc.; 1999. p. 293–308.

38. Faas FG, Avramut MC, van den Berg BM, Mommaas AM, Koster AJ, Ravelli RB. Virtual nanoscopy: generation of ultra-large high resolution electron microscopy maps. J Cell Biol. 2012;198(3):457–69. Epub 2012/08/08. doi: 10.1083/jcb.201201140. PubMed PMID: 22869601; PubMed Central PMCID: PMCPMC3413355.

39. Mayhew TM, Lucocq JM, Griffiths G. Relative labelling index: a novel stereological approach to test for non-random immunogold labelling of organelles and membranes on transmission electron microscopy thin sections. Journal of microscopy. 2002;205(Pt 2):153–64. PubMed PMID: 11879430.

40. Tooze J, Tooze S, Warren G. Replication of coronavirus MHV-A59 in sac-cells: determination of the first site of budding of progeny virions. Eur J Cell Biol. 1984;33(2):281–93. Epub 1984/03/01. PubMed PMID: 6325194.

41. Goldsmith CS, Tatti KM, Ksiazek TG, Rollin PE, Comer JA, Lee WW, et al. Ultrastructural characterization of SARS coronavirus. Emerg Infect Dis. 2004;10(2):320–6. Epub 2004/03/20. doi: 10.3201/eid1002.030913. PubMed PMID: 15030705; PubMed Central PMCID: PMCPMC3322934.

42. Stertz S, Reichelt M, Spiegel M, Kuri T, Martinez-Sobrido L, Garcia-Sastre A, et al. The intracellular sites of early replication and budding of SARS-coronavirus. Virology. 2007;361(2):304–15. Epub 2007/01/11. doi: 10.1016/j.virol.2006.11.027. PubMed PMID: 17210170.

43. Karreman MA, Van Donselaar EG, Agronskaia AV, Verrips CT, Gerritsen HC. Novel contrasting and labeling procedures for correlative microscopy of thawed cryosections. J Histochem Cytochem. 2013;61(3):236–47. Epub 2012/12/25. doi: 10.1369/0022155412473756. PubMed PMID: 23264637; PubMed Central PMCID: PMCPMC3636698.

44. Klumperman J, Locker JK, Meijer A, Horzinek MC, Geuze HJ, Rottier PJ. Coronavirus M proteins accumulate in the Golgi complex beyond the site of virion budding. J Virol. 1994;68(10):6523–34. Epub 1994/10/01. PubMed PMID: 8083990; PubMed Central PMCID: PMCPMC237073.

45. Ng ML, Tan SH, See EE, Ooi EE, Ling AE. Proliferative growth of SARS coronavirus in Vero E6 cells. J Gen Virol. 2003;84(Pt 12):3291–303. doi: 10.1099/vir.0.19505-0. PubMed PMID: 14645910.

46. Becker WB, McIntosh K, Dees JH, Chanock RM. Morphogenesis of avian infectious bronchitis virus and a related human virus (strain 229E). J Virol. 1967;1(5):1019–27. Epub 1967/10/01. PubMed PMID: 5630226; PubMed Central PMCID: PMCPMC375381.

47. Zhou X, Cong Y, Veenendaal T, Klumperman J, Shi D, Mari M, et al. Ultrastructural Characterization of Membrane Rearrangements Induced by Porcine Epidemic Diarrhea Virus Infection. Viruses. 2017;9(9). Epub 2017/09/06. doi: 10.3390/v9090251. PubMed PMID: 28872588; PubMed Central PMCID: PMCPMC5618017.

48. Machamer CE, Mentone SA, Rose JK, Farquhar MG. The E1 glycoprotein of an avian coronavirus is targeted to the cis Golgi complex. Proc Natl Acad Sci U S A. 1990;87(18):6944–8. Epub 1990/09/01. doi: 10.1073/pnas.87.18.6944. PubMed PMID: 2169615; PubMed Central PMCID: PMCPMC54658.

49. Opstelten DJ, Raamsman MJ, Wolfs K, Horzinek MC, Rottier PJ. Envelope glycoprotein interactions in coronavirus assembly. J Cell Biol. 1995;131(2):339–49. Epub 1995/10/01. doi: 10.1083/jcb.131.2.339. PubMed PMID: 7593163; PubMed Central PMCID: PMCPMC2199982.

50. Denison MR, Spaan WJ, van der Meer Y, Gibson CA, Sims AC, Prentice E, et al. The putative helicase of the coronavirus mouse hepatitis virus is processed from the replicase gene polyprotein and localizes in complexes that are active in viral RNA synthesis. J Virol. 1999;73(8):6862–71. Epub 1999/07/10. PubMed PMID: 10400784; PubMed Central PMCID: PMCPMC112771.

51. van der Meer Y, Snijder EJ, Dobbe JC, Schleich S, Denison MR, Spaan WJ, et al. Localization of mouse hepatitis virus nonstructural proteins and RNA synthesis indicates a role for late endosomes in viral replication. J Virol. 1999;73(9):7641–57. Epub 1999/08/10. PubMed PMID: 10438855; PubMed Central PMCID: PMCPMC104292.

52. Sims AC, Ostermann J, Denison MR. Mouse hepatitis virus replicase proteins associate with two distinct populations of intracellular membranes. J Virol. 2000;74(12):5647–54. Epub 2000/05/24. doi: 10.1128/jvi.74.12.5647-5654.2000. PubMed PMID: 10823872; PubMed Central PMCID: PMCPMC112052.

53. V’Kovski P, Gerber M, Kelly J, Pfaender S, Ebert N, Braga Lagache S, et al. Determination of host proteins composing the microenvironment of coronavirus replicase complexes by proximity-labeling. eLife. 2019;8. doi: 10.7554/eLife.42037. PubMed PMID: 30632963; PubMed Central PMCID: PMC6372286.

54. McBride R, van Zyl M, Fielding BC. The coronavirus nucleocapsid is a multifunctional protein. Viruses. 2014;6(8):2991–3018. Epub 2014/08/12. doi: 10.3390/v6082991. PubMed PMID: 25105276; PubMed Central PMCID: PMCPMC4147684.

55. Hagemeijer MC, Vonk AM, Monastyrska I, Rottier PJ, de Haan CA. Visualizing coronavirus RNA synthesis in time by using click chemistry. J Virol. 2012;86(10):5808–16. doi: 10.1128/JVI.07207-11. PubMed PMID: 22438542; PubMed Central PMCID: PMC3347275.

56. Deng Y, Almsherqi ZA, Ng MM, Kohlwein SD. Do viruses subvert cholesterol homeostasis to induce host cubic membranes? Trends in cell biology. 2010;20(7):371–9. doi: 10.1016/j.tcb.2010.04.001. PubMed PMID: 20434915.

57. Almsherqi ZA, Kohlwein SD, Deng Y. Cubic membranes: a legend beyond the Flatland* of cell membrane organization. J Cell Biol. 2006;173(6):839–44. doi: 10.1083/jcb.200603055. PubMed PMID: 16785319; PubMed Central PMCID: PMCPMC2063909.

58. Paul D, Hoppe S, Saher G, Krijnse-Locker J, Bartenschlager R. Morphological and biochemical characterization of the membranous hepatitis C virus replication compartment. J Virol. 2013;87(19):10612–27. Epub 2013/07/26. doi: 10.1128/JVI.01370-13. PubMed PMID: 23885072; PubMed Central PMCID: PMCPMC3807400.

59. Melia CE, van der Schaar HM, Lyoo H, Limpens R, Feng Q, Wahedi M, et al. Escaping Host Factor PI4KB Inhibition: Enterovirus Genomic RNA Replication in the Absence of Replication Organelles. Cell reports. 2017;21(3):587–99. doi: 10.1016/j.celrep.2017.09.068. PubMed PMID: 29045829.

60. Melia CE, van der Schaar HM, de Jong AWM, Lyoo HR, Snijder EJ, Koster AJ, et al. The Origin, Dynamic Morphology, and PI4P-Independent Formation of Encephalomyocarditis Virus Replication Organelles. mBio. 2018;9(2). Epub 2018/04/19. doi: 10.1128/mBio.00420-18. PubMed PMID: 29666283; PubMed Central PMCID: PMCPMC5904412.

61. van Hemert MJ, van den Worm SH, Knoops K, Mommaas AM, Gorbalenya AE, Snijder EJ. SARS-coronavirus replication/transcription complexes are membrane-protected and need a host factor for activity in vitro. PLoS Pathog. 2008;4(5):e1000054. Epub 2008/05/03. doi: 10.1371/journal.ppat.1000054. PubMed PMID: 18451981; PubMed Central PMCID: PMCPMC2322833.

62. Kindler E, Gil-Cruz C, Spanier J, Li Y, Wilhelm J, Rabouw HH, et al. Early endonuclease-mediated evasion of RNA sensing ensures efficient coronavirus replication. PLoS Pathog. 2017;13(2):e1006195. Epub 2017/02/06. doi: 10.1371/journal.ppat.1006195. PubMed PMID: 28158275; PubMed Central PMCID: PMCPMC5310923.

63. Thiel V, Ivanov KA, Putics A, Hertzig T, Schelle B, Bayer S, et al. Mechanisms and enzymes involved in SARS coronavirus genome expression. J Gen Virol. 2003;84(Pt 9):2305–15. Epub 2003/08/15. doi: 10.1099/vir.0.19424-0. PubMed PMID: 12917450.

64. Hamre D, Procknow JJ. A new virus isolated from the human respiratory tract. Proc Soc Exp Biol Med. 1966;121(1):190–3. Epub 1966/01/01. doi: 10.3181/00379727-121-30734. PubMed PMID: 4285768.

65. Casais R, Thiel V, Siddell SG, Cavanagh D, Britton P. Reverse genetics system for the avian coronavirus infectious bronchitis virus. J Virol. 2001;75(24):12359–69. Epub 2001/11/17. doi: 10.1128/JVI.75.24.12359-12369.2001. PubMed PMID: 11711626; PubMed Central PMCID: PMCPMC116132.

66. Bosch BJ, van der Zee R, de Haan CA, Rottier PJ. The coronavirus spike protein is a class I virus fusion protein: structural and functional characterization of the fusion core complex. J Virol. 2003;77(16):8801–11. doi: 10.1128/jvi.77.16.8801-8811.2003. PubMed PMID: 12885899; PubMed Central PMCID: PMC167208.

67. Snijder EJ, van der Meer Y, Zevenhoven-Dobbe J, Onderwater JJ, van der Meulen J, Koerten HK, et al. Ultrastructure and origin of membrane vesicles associated with the severe acute respiratory syndrome coronavirus replication complex. J Virol. 2006;80(12):5927–40. doi: 10.1128/JVI.02501-05. PubMed PMID: 16731931; PubMed Central PMCID: PMC1472606.

68. Schonborn J, Oberstrass J, Breyel E, Tittgen J, Schumacher J, Lukacs N. Monoclonal antibodies to double-stranded RNA as probes of RNA structure in crude nucleic acid extracts. Nucleic acids research. 1991;19(11):2993–3000. doi: 10.1093/nar/19.11.2993. PubMed PMID: 2057357; PubMed Central PMCID: PMC328262.

69. Widjaja I, Wang C, van Haperen R, Gutierrez-Alvarez J, van Dieren B, Okba NMA, et al. Towards a solution to MERS: protective human monoclonal antibodies targeting different domains and functions of the MERS-coronavirus spike glycoprotein. Emerging microbes & infections. 2019;8(1):516–30. doi: 10.1080/22221751.2019.1597644. PubMed PMID: 30938227; PubMed Central PMCID: PMC6455120.

70. Ginsel LA, Onderwater JJ, Daems WT. Resolution of a gold latensification-elon ascorbic acid developer for Ilford L4 emulsion. Histochemistry. 1979;61(3):343–6. PubMed PMID: 478993.

71. Kukulski W, Schorb M, Welsch S, Picco A, Kaksonen M, Briggs JA. Correlated fluorescence and 3D electron microscopy with high sensitivity and spatial precision. J Cell Biol. 2011;192(1):111–9. doi: 10.1083/jcb.201009037. PubMed PMID: 21200030; PubMed Central PMCID: PMC3019550.

72. Kremer JR, Mastronarde DN, McIntosh JR. Computer visualization of three-dimensional image data using IMOD. Journal of structural biology. 1996;116(1):71–6. doi: 10.1006/jsbi.1996.0013. PubMed PMID: 8742726.

73. Lucocq JM, Habermann A, Watt S, Backer JM, Mayhew TM, Griffiths G. A rapid method for assessing the distribution of gold labeling on thin sections. J Histochem Cytochem. 2004;52(8):991–1000. doi: 10.1369/jhc.3A6178.2004. PubMed PMID: 15258174.

## References Supporting Information

1. Bienz KA. Techniques and applications of autoradiography in the light and electron microscope. Microsc Acta. 1977;79(1):1–22. Epub 1977/01/01. PubMed PMID: 65723.

2. Bozzola JJ, Russell LD. Autoradiography & Radioautography. Electron Microscopy: Principles and Techniques for Biologists. Sudbury, MA.: Jones and Bartlett Publishers, Inc.; 1999. p. 293–308.

3. Bienz K, Egger D, Pasamontes L. Association of polioviral proteins of the P2 genomic region with the viral replication complex and virus-induced membrane synthesis as visualized by electron microscopic immunocytochemistry and autoradiography. Virology. 1987;160(1):220–6. PubMed PMID: 2820130.

4. Melia CE, van der Schaar HM, Lyoo H, Limpens R, Feng Q, Wahedi M, et al. Escaping Host Factor PI4KB Inhibition: Enterovirus Genomic RNA Replication in the Absence of Replication Organelles. Cell reports. 2017;21(3):587–99. doi: 10.1016/j.celrep.2017.09.068. PubMed PMID: 29045829.

5. Melia CE, van der Schaar HM, de Jong AWM, Lyoo HR, Snijder EJ, Koster AJ, et al. The Origin, Dynamic Morphology, and PI4P-Independent Formation of Encephalomyocarditis Virus Replication Organelles. mBio. 2018;9(2). Epub 2018/04/19. doi: 10.1128/mBio.00420-18. PubMed PMID: 29666283; PubMed Central PMCID: PMCPMC5904412.

6. de Wilde AH, Raj VS, Oudshoorn D, Bestebroer TM, van Nieuwkoop S, Limpens RW, et al. MERS-coronavirus replication induces severe in vitro cytopathology and is strongly inhibited by cyclosporin A or interferon-alpha treatment. J Gen Virol. 2013;94(Pt 8):1749–60. Epub 2013/04/27. doi: 10.1099/vir.0.052910-0. PubMed PMID: 23620378; PubMed Central PMCID: PMCPMC3749523.

7. Knoops K, Kikkert M, Worm SH, Zevenhoven-Dobbe JC, van der Meer Y, Koster AJ, et al. SARS-coronavirus replication is supported by a reticulovesicular network of modified endoplasmic reticulum. PLoS biology. 2008;6(9):e226. doi: 10.1371/journal.pbio.0060226. PubMed PMID: 18798692; PubMed Central PMCID: PMC2535663.

